# Bumblebee worker body size affects new worker production in different resource environments

**DOI:** 10.1101/2020.01.16.909135

**Authors:** Natalie Z. Kerr, Rosemary L. Malfi, Neal M. Williams, Elizabeth E. Crone

**Affiliations:** Department of Biology, Tufts University, Medford, Massachusetts 02155, USA; Department of Biology, Duke University, Durham, North Carolina 27710, USA; Department of Biology, University of Massachusetts-Amherst, Amherst, MA 01003, USA; Department of Entomology and Nematology, University of California, Davis, CA 95616, USA

**Keywords:** *Bombus vosnesenskii*, functional linear models, colony age, egg production, larval survival, development, callow size

## Abstract

1. Behavior and organization of social groups is thought to be vital to the functioning of societies, yet the contributions of various roles within social groups have been difficult to quantify. A common approach to quantifying these role-based contributions is evaluating the performance of individuals at conducting certain roles, these studies ignore how these performances might scale up to effects at the population-level. Manipulative experiments are another common approach to determine population-level effects, but they often ignore potential feedbacks associated with these various roles.
2. Here, we evaluate the effects of worker size distribution in bumblebee colonies on worker production, using functional linear models. Functional linear models are a recent correlative technique that has been used to assess lag effects of environmental drivers on plant performance. We demonstrate potential applications of this technique to explore contributions of social animals to ecological phenomenon.
3. We found that the worker size distribution differentially affected new worker production across three resource environments. Specifically, more larger workers had mostly positive effects and more smaller workers had negative effects on worker production. Most of these effects were only detected under low or fluctuating resource environments suggesting that the advantage of colonies with larger-bodied workers becomes more apparent under stressful conditions.
4. We demonstrate the wider ecological application of functional linear models. We highlight the advantages and limitations when considering these models, and how they are a valuable complement to many of these performance-based and manipulative experiments.

## INTRODUCTION

In animal societies, individuals are often observed performing different tasks, such as guarding nests and burrows (Clutton-Brock *et al*. 2001a), nursing and caring for young (Wilkinson 1992; Kerth 2008; Sparkman *et al*. 2011), or reproducing (Jarvis 1981; Faulkes & Bennett 2001). The roles within these social groups are commonly assigned based on the age (Jarvis 1981; Seeley & Kolmes 1991; Brent *et al*. 2015; Zöttl *et al*. 2016), size (Porter & Tschinkel 1985; Wenzel 1992; Schwander, Rosset & Chapuisat 2005; Goulson 2009), and/or status (Frank 1986; Sparkman *et al*. 2011) of individuals. For example, in Meerkats, which are cooperative breeders, younger non-breeding individuals often stand on ‘sentinel duty’ during group foraging bouts and care for offspring of the dominant breeding pair (Clutton-Brock *et al*. 2001b; Clutton-Brock *et al*. 2002; Clutton-Brock, Russell & Sharpe 2004). Without the co-operation of these non-breeders, the survival of individuals within the colonies is likely to decrease, particularly for the young (Doolan & Macdonald 1999; Russell *et al*. 2007). This social behavior and organization is often assumed to be vital to the functioning and survival of these societies.

The most common approach to understanding the contribution of roles within social groups is to observe the behavior and performance of individuals. However, most observational performance-based investigations do not quantify the contribution of individuals that perform certain roles to the functioning of the society. To attempt to tackle the challenges associated with quantifying trait-based contributions, a few studies have manipulated colonies in the laboratory to evaluate the effects of the social organization of age- and size-polymorphic species, such as mole rats (Jarvis 1981; Zöttl *et al*. 2016), ants (Porter & Tschinkel 1985; Billick & Carter 2007), and bumblebees (Cnaani & Hefetz 1994; Jandt & Dornhaus 2009; Couvillon *et al*. 2010; Jandt & Dornhaus 2011; Jandt & Dornhaus 2014). In laboratory colonies of a eusocial ant *Pheidole dentata*, larvae gained more mass when reared by older workers, suggesting that older workers contribute more towards worker production in these ant colonies than their younger sisters (Muscedere, Willey & Traniello 2009). However, colonies within these laboratory experiments were not faced with the same external environmental stressors as those in the wild. In the case of bumblebees, larger workers are more susceptible to predators and parasites (Cartar & Dill 1991; Muller, Blackburn & Schmid-Hempel 1996; Malfi & Roulston 2014), despite being better foragers. Therefore, the behaviors of social organism under artificial conditions might not capture all the feedbacks associated with size or age-based roles.

A recent statistical approach called functional linear models (FLMs) provides an alternative method of inference to manipulative laboratory experiments. FLMs can evaluate the contributions of age- or size-based roles within societies using observational data. FLMs assume that the effect of a predictor variable (e.g. number of workers) on a response variable (e.g. egg production) is a smooth function of some feature of the predictor variable (e.g. size of workers). Past applications of FLMs in ecology have investigated environmental drivers of plant population dynamics (Teller *et al*. 2016; Tenhumberg *et al*. 2018). These studies evaluated the effects of environmental conditions (e.g. precipitation) on plant performance (e.g. growth) assuming the slope of the effect of environmental conditions and plant performance varies as a smooth function of the time lag between conditions and performance (e.g. precipitation in the past 1, 2, 3… months). For example, the slope of precipitation versus plant growth could go from positive in recent months to zero at longer time lags. This method has potential for wider ecological application to investigate life history phenomena. Here, we explore application of FLMs to quantifying the relationship between aspects of new worker production as a function of the body size of existing workers in bumblebee colonies.

Bumblebees (*Bombus* spp.) are primitively eusocial insects that form relatively small colonies and have a discrete life cycle lasting only for a single season, which makes them a tractable system for studying trait-based roles within societies. Bumblebees also exhibit worker size polymorphism, where workers within colonies vary up to 10-fold in mass (Goulson 2009). In bumblebee colonies, larger workers are often found foraging and guarding, while smaller workers spend more time in the colony conducting in-nest tasks such as fanning and incubating (Richards 1946; Cumber 1949; Goulson *et al*. 2002; Jandt & Dornhaus 2009; Inoue *et al*. 2010). Many studies have measured the importance of body size in determining how workers perform various tasks, ranging from foraging and flight dynamics to thermoregulating and undertaking. Most of these have found that larger workers are better at multiple tasks, such as foraging and nursing (Cnaani & Hefetz 1994; Goulson *et al*. 2002; Spaethe & Weidenmüller 2002; Peat & Goulson 2005; Ings 2007; Spaethe *et al*. 2007; Kerr, Crone & Williams 2019), with a few studies concluding either that intermediate-size is better (Jandt & Dornhaus 2014), or that there is no size-based difference (Jandt & Dornhaus 2014). Although these studies demonstrate that body size affects worker performance at certain tasks, they do not demonstrate how their size-based performance at tasks may, in turn, affect colony growth and development.

No studies have found smaller bumblebee workers to be better at performing tasks essential to colony function. However, smaller workers are more resilient to starvation (Couvillon & Dornhaus 2010). Therefore, their value may become more apparent when food resources are limiting. In addition, smaller workers have lower production costs, so may be more cost-effective (Kerr, Crone & Williams 2019). Here, we used FLMs to evaluate the contribution of worker size polymorphism to worker production in bumblebee colonies under three different resource environments: a low resource environment; an environment with an early season pulse followed by low resources (‘high-low’); and a high resource environment. We looked at five vital rates relating to worker production: (1) number of new eggs laid, (2) development time, (3) larval survival, and (4) mean and (5) variance in worker emergence size, i.e. the size of callow workers. By evaluating the contribution of different-sized workers under different resources environments to worker production, we can assess whether larger workers are more beneficial when resource conditions are more favorable and whether the benefit of small workers to colonies is only seen when resources are low, making both production cost and resistance to starvation a premium.

## MATERIALS AND METHODS

### Study species and sites

We hand reared *Bombus vosnesenskii* colonies from wild-caught queens collected at the University of California McLaughlin Reserve (N38 52 25.74, W122 25 56.25) in early spring 2015 and 2016 while they searched for nest sites. These colonies are the same as used for other studies (Kerr, Crone & Williams 2019; Malfi, Crone & Williams 2019), so we only briefly describe the rearing process here.

In 2015 and 2016, we hand-reared colonies in the laboratory in a dark room at 26-28°C for 6 to 9 weeks until their second or first cohort of worker bees eclosed. In 2015, we relocated seven colonies outside (N38 32 12.21, W121 47 16.95) at the Harry H. Laidlaw Jr. Honey Bee Research Facility (Davis, CA), where the surrounding landscape consisted of agricultural crops, floral research plots, and a 0.2 ha pollinator garden (Fig. S1a). In 2016, we relocated 14 colonies outside in agricultural fields at UC Davis Experimental Farm property (N38 31 32.3, W121 46 56.54). Half of the colonies (n = 7) had access to flight cages that provided a pulse of native California wildflower species for ∼4 weeks early in the season (“pulse” treatment) and the other half had no supplemental forage (“control” treatment) (Malfi, Crone & Williams 2019). The surrounding landscapes were croplands consisting of mainly non-flowering cereals, corn, and a strip of riparian habitat (Fig. S1b).

In this study, we broadly categorized the resource environments experienced by our experimental colonies in each of these years according to the amount of available forage. The 2015 colonies, located next to a pollinator garden at the Honey Bee Research Facility, had the highest resource availability (“high”), colonies in the 2016 pulse treatment had the second highest resource availability (“high-low”), and colonies in the 2016 control treatment had the lowest availability (“low”). These three environments will now be referred to as high, high-low, and low.

### Brood mapping

Each week, we photographed the brood from multiple angles (aerial, side, diagonal) to fully capture all brood cells. We individually numbered each brood cell as it differentiated and tracked the fate of all marked cells throughout colony development (Fig. 1). We classified each living brood cell into five categories: (1) clump stage, which represents the egg stage where individual cells have not yet differentiated; (2) pre-differentiated stage, which represents early larval instars where individual cells have begun differentiating; (3) differentiated stage, which represents later larval instars where individual brood cells are clearly differentiated; (4) cocoon stage, where cells have darkened indicating that pupa have spun their cocoons; and (5) eclosed stage, where cell has opened and an adult worker emerged (Fig. 2 for stages). We also had two other categories: (6) dead, where we have observed a dead cell, and (7) unseen, where the cell can no longer be seen in the brood photos.

**Figure 1.**
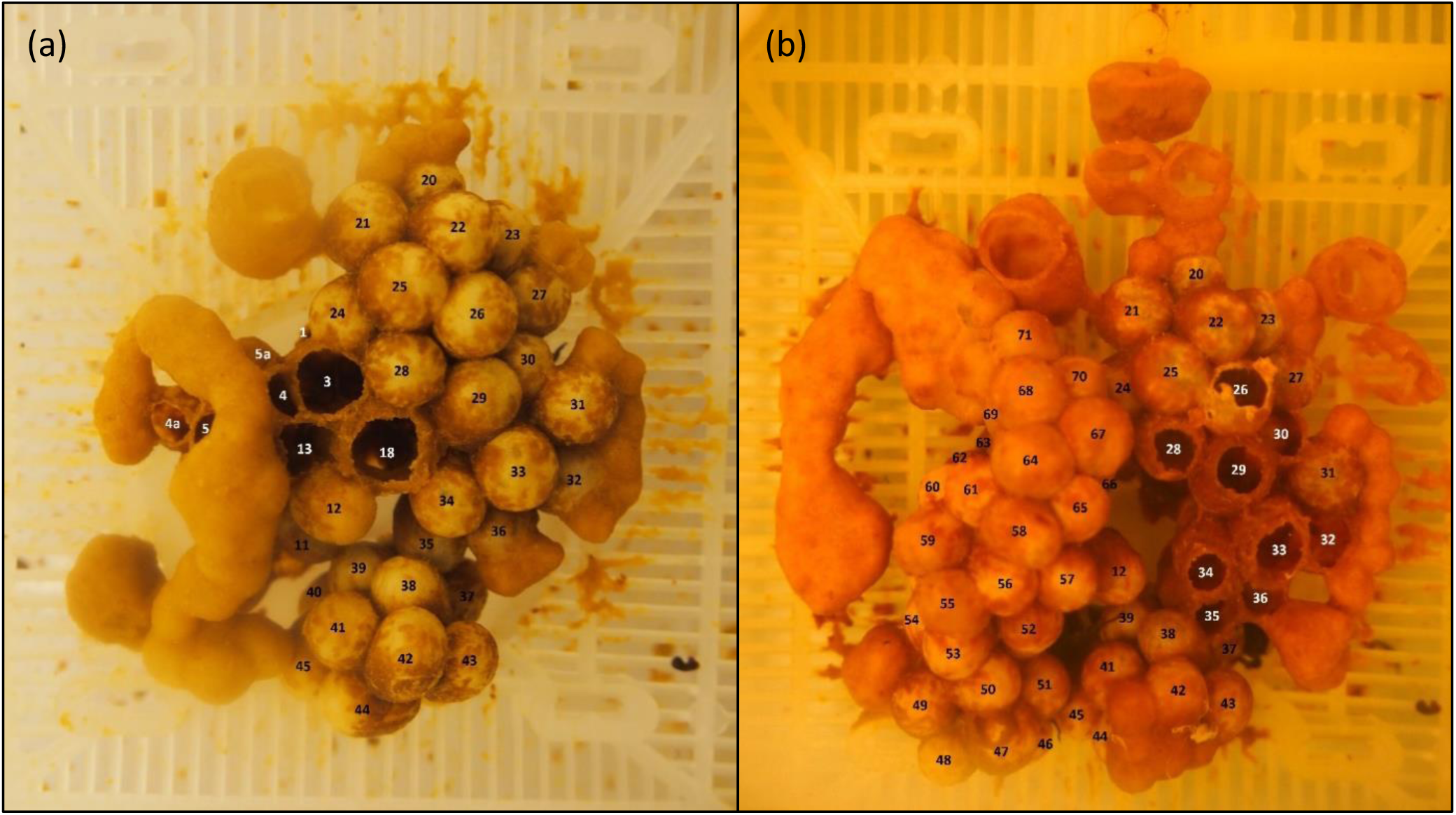
Example of brood mapping photos used to track the fate of individual cells. These mapping photos are aerial photographs for colony 6 in (a) week 5 and (b) week 6 since the first brood photo. Aerial, side, and diagonal photos were taken to capture all cells. Each cell has been individually numbered to track each cell.

**Figure 2.**
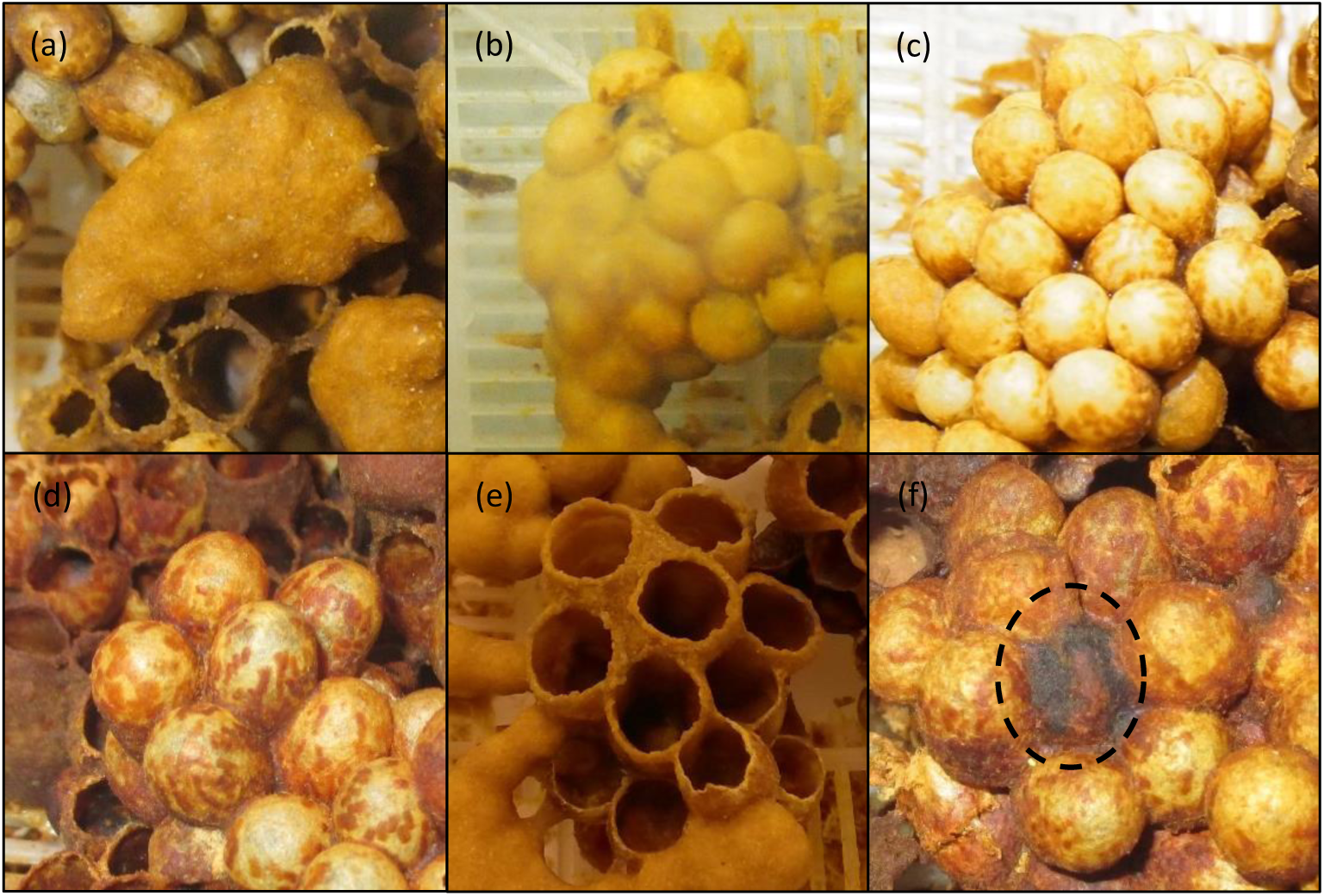
Brood mapping photos showing each of the six categories of living or dead stages of cell development. The six stages are: (a) clump stage, which are egg stages; (b) pre-popcorn stages, which represents early larval instars; (c) popcorn stage, which are late instar larvae; (d) cocoon stage; (e) eclosed stage, and (f) a dead cell (dashed circle). These categories assisted with estimating three vital rates: eggs laid, development time, and larval survival.

Some brood clumps did not develop into distinct cells before the end of brood mapping, while other clumps died before cell partitioning. Rather than exclude these indistinct, dead, or undeveloped brood clumps in our analyses, which could result in underestimating egg production and overestimating larval survival, we estimated the number of cells for these clumps. We did this by classifying these indistinct brood clumps into five size categories (tiny, small, medium, large, extra-large) based on comparisons with similarly-sized brood clumps that did divide into individual cells and assigning the mean value of cells for these size categories to indistinct clumps.

From the brood mapping, we estimated three vital rates: egg production, larval development time, and larval survival. We considered weekly egg production to be the number of newly visible cells in either clump or pre-differentiated stages. We assumed that the number of distinct cells formed by a brood clump represented the total number of eggs laid, i.e. no eggs died before larval cells differentiated. We calculated development time for each cell as the number of days from when it was first seen as an egg (defined as the ‘clump’ stage) to when it was first seen as an eclosed cell. Cells that were not detected in the clump stage or that disappeared from view before visibly eclosing were excluded from our analyses of larval development time. Finally, we classified larval survival as the success of each cell reaching eclosion. We excluded 43 unseen brood cells from our larval analyses because more than 8 days (50% the normal bumblebee development time) passed between photos of them so their fates could not be unambiguously mapped. These represent 10% of 437 unseen cells or 1% of all 4,640 cells mapped across the 21 colonies and three resource environments.

### Worker surveys

We conducted weekly night-time surveys to estimate the mean and standard deviation in the size of newly emerged workers (hereafter referred to as “callow size”). We assigned each bee a unique tag using a combination enamel paint and numbered, color-tags or Microsensys radio-frequency identification (RFID) tags (Kerr, Crone & Williams 2019; Malfi, Crone & Williams 2019). For each newly emerged (“callow”) worker, we estimated body size by measuring intertegular (IT) span to the nearest 0.01 mm using digital calipers (Cane 1987; Hagen & Dupont 2013) and wet weight to the nearest 0.01 mg using analytical microbalance (Mettler Toledo XS205DU). The size of each worker at initial capture was used to estimate the mean and standard deviation in callow size. We used these size measurements in combination with presence/absence data to determine the number of workers of each size (now referred to as “worker size composition”) present in each colony for each week of the survey in order to evaluate the effects of worker size composition on aspects of worker production.

### Functional linear models

We used functional linear models (FLMs) to estimate how five vital rates varied with worker size composition. FLMs are a type of regression spline that allows a covariate to vary smoothly over a continuous domain (Ramsay & Silverman 2005; Ramsay, Hooker & Graves 2009). Therefore, instead of restricting our predictors (X) to unidimensional space (i.e. simple linear models, such as total worker number predicts number of eggs), we can evaluate the effect of the number of workers on some response variable (e.g. number of eggs) as a continuous function of worker size (i.e. a separate attribute of the predictor variable), such that the smooth function of size-specific slopes versus worker size can be described as:

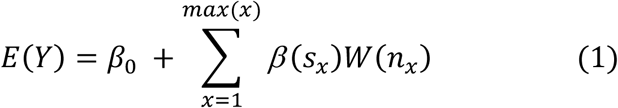

where *E*(*Y*) is the expected value of the response variable *Y* (e.g. number of eggs); *β*_0_ is the intercept; *W*(*n*_*x*_) is the number of workers *n* of size *x*; and *β (s)* is the slope of *Y* versus the number of workers of each size category *x* (c.f. methods in Teller *et al*. 2016). Here, the continuous attribute (i.e. worker size) of the predictor variable (i.e. number of workers) is discretized into many size categories (14 size categories for both low and high-low, and 17 for high resource colonies) to approximate a continuous distribution of sizes (i.e., the worker size composition). The expected value of the response variable is the sum of the product of the size-specific slopes *β (s*_*x*_) multiplied by the number of workers of size *x* (Fig. 3). If the slope of *Y* versus the number of workers of size *x* is positive, then more workers of size *x* increase values of *Y* and vice versa when the slope is negative (Fig. 3).

**Figure 3.**
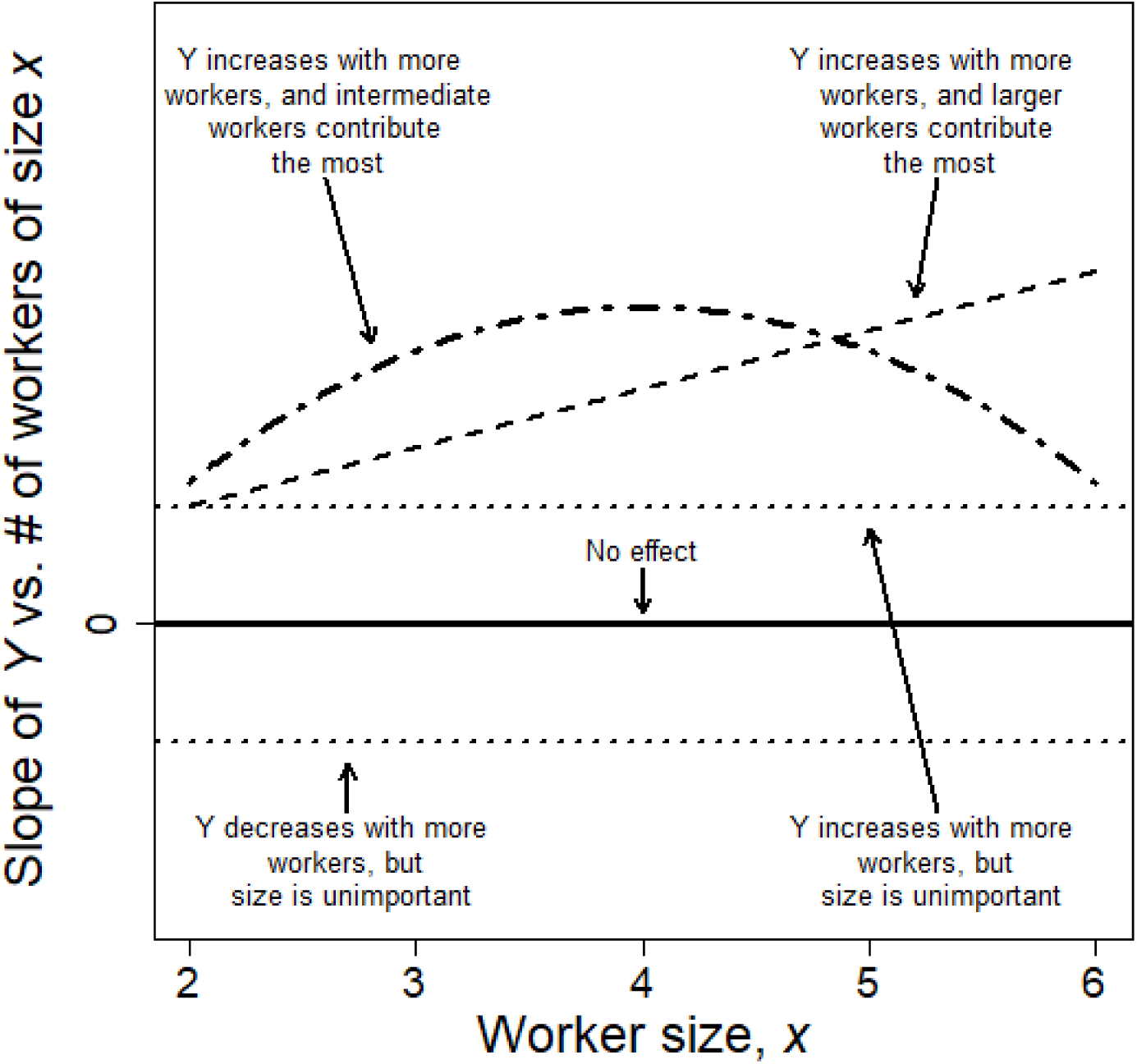
Example of functional linear model results showing the smooth function of the slopes of Y versus the number of workers of size *x* as a function of worker size, *x. Y* covariate could be one of the five metrics of worker production: egg production, larval development time, larval survival, and mean and variance in callow size. We illustrate the following examples, where # number of workers or their size has no effect on *Y* (solid line); either more workers increases (β_0_ > 0) or decreases (β_0_ < 0) *Y*, but size is unimportant (dotted lines); more workers increase *Y* and larger workers contribute the most (dashed line); and more increase *Y* and intermediate workers contribute the most (dotted and dashed line).

We parameterized the smooth functions of the size-specific slopes using general additive models (GAMs). We fit our GAMs using the cubic spline basis for all smooth covariates, so that the coefficients will be set to 0 if our covariates have no effects on the response (see Zuur 2012, for an excellent textbook introduction to GAMs). For new eggs laid, we used worker size composition in the previous week to predict the number of eggs laid in the present time step for our size composition FLMs. For the other three vital rates, we quantified worker size composition as the average number of workers in each size category across their larval development period.

Models were fit separately to data from each study (i.e. low, high-low, and high resource environments), and we included colony ID as a fixed effect (i.e. a different intercept term for each colony) for each model. We used negative binomial GAMs to account for overdispersion for estimating new eggs laid and development time. We offset the number of new eggs laid by the number of days between brood photos. We used binomial and Gaussian-distributed GAMs for larval survival and callow size, respectively. We parameterized the binomial GAMs for estimating larval survival using successes and failures, where the total number of trials was defined as the number of days between brood photos, and the number of failures was defined as “1” if the cell died and “0” if it survived. We restricted the number of knots for our smooth terms of the number of workers of size *j* to a maximum of five. We also rejected any model structure that did not produce unimodal functions for our smooth term of worker size composition, since GAMs are prone to overfitting, and multimodal functions generally did not appear to be biologically meaningful. We used likelihood ratio tests to assess the fit of the parametric intercept term and the number of knots for each smooth term in our models given our data. We ran these general additive models (using mgcv::gam; Wood 2004; Wood 2011) in program R (R Core Team 2017). We repeated all analyses with slopes scaled to worker production costs, rather than numbers of individuals. Because these results were largely parallel (Appendix S2), we do not discuss them further.

Colony size (i.e. number of observed workers) increased with colony age across three resource environments (Fig. S2-4). To avoid potentially confounding effects due to collinearity between colony age and worker number, we ran models separately with colony age and worker size composition as predictors of various measures of worker production success. Results for colony age are described in Appendix S1. Relationships between worker survival and larval survival and mean callow size in the low resource environment were somewhat confounded with colony age effects, and should be interpreted with caution (Table 1, Appendix S4). We found no evidence for other pairwise relationships for colony age, worker number, mean worker size, and standard deviation in worker size across the three resource environments.

**Table 1.** Relationship of the five vital rates relating to worker production and the smooth terms of colony age, the number of workers of each size (i.e. worker size composition, WSC), and standardized (“std”) WSC meaning that we accounted for production costs associated with each worker size. Relationship descriptions provided are restricted over the observed range of worker body sizes and colony ages including days spent in the laboratory. The column “confounding effects” describes whether both colony age and WSC had similar effects on the response variable when both smooth terms are significant. Since colony age and population size are correlated, we were unable to determined which smooth term was driving these effects if both smooth terms have similar effects. Asterisks (*) represents a significant effect of a main effect of colony ID on the parametric intercept in the GAMs.

| Response variable | Resource environment | Smooth terms |  |  | Confounding effects |
| --- | --- | --- | --- | --- | --- |
|  |  | *Colony age* | *WSC* | *Std WSC* |  |
| Egg production | *Low* | Concave* | – * | – | – |
|  | *High-low* | Concave* | Increases* | Increases | Possibly |
|  | *High* | – * | Increases | Increases | – |
| Development time | *Low* | Multimodal* | Decreases* | Decreases | Possibly |
|  | *High-low* | Concave* | Decreases* | Decreases | Possibly |
|  | *High* | – * | – * | Decreases | – |
| Larval survival | *Low* | Decreases* | Increases* | Increases | Yes |
|  | *High-low* | Decreases* | Increases* | Increases | Yes |
|  | *High* | Decreases* | Decreases* | Decreases | No |
| Mean callow size | *Low* | Decrease | Increases | Increases | Yes |
|  | *High-low* | Multimodal* | – * | Decreases | – |
|  | *High* | Decreases* | Decreases* | – | No |
| Standard deviation in callow size | *Low* | – | – | – | – |
|  | *High-low* | – | Decreases | Decreases | – |
|  | *High* | – | – | Decreases | – |

## RESULTS

Average worker size increased with available ambient resources. Malfi et al (2019) report smaller worker size in the low (mean and SE in IT span: 3.19 ± 0.03) than the high-low (IT span: 3.32 ± 0.06) resource environment. Consistent with this pattern, workers were largest in the high resource environments (IT span: 3.68 ± 0.04).

### Daily egg production

Worker size composition did not affect egg production in the low resource environment (Fig. 4a; low – *χ*^*2*^ = 1E-4, d.f = 5.1E-5, *P* = 0.15). More larger workers increased egg production in both the high-low and high resource environments (Fig. 4b-c; high-low – *χ*^*2*^ = 11.5, d.f. = 1.1, *P* < 0.001; high - *χ*^*2*^ = 5.6, d.f. = 1.1, *P* = 0.01), but more larger workers had greater impact on egg production in the high-low resource environment than in the constantly high resource environment.

**Figure 4.**
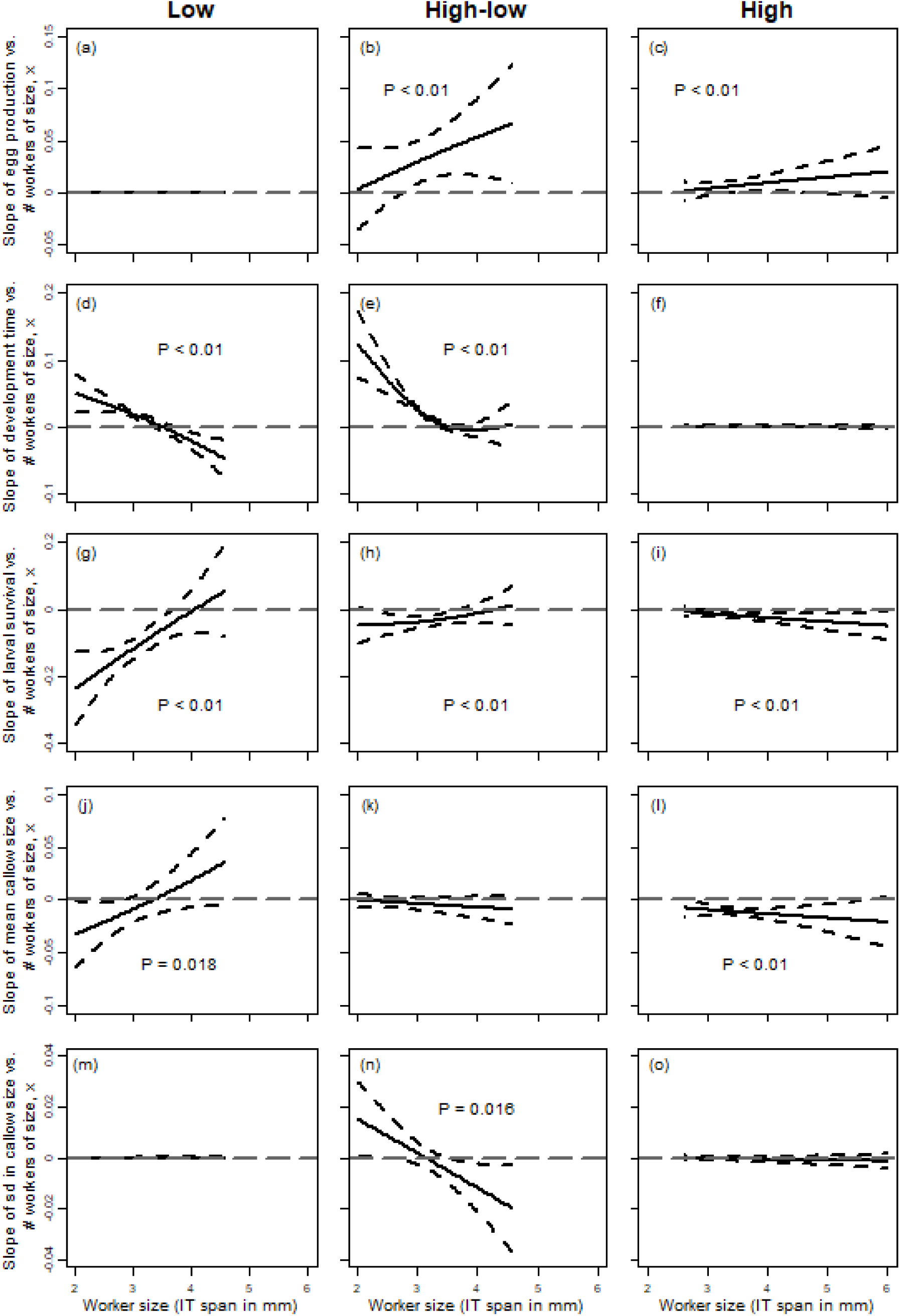
Generalized additive model results depicting the smooth function of the slopes for all five vital rates versus the number of workers of size x as a function of worker size x for the low (left), high-low (middle), and high (right) resource environments. Worker size was measured as the distance between the tegulae, i.e. intertegular (IT) span in mm. Grey dotted horizontal line at zero represent deviations from mean slope values, i.e. slopes above the line means more workers of size x have positive impact on Y. Plots with a significant smooth term of WSC are labeled with P < 0.01. Note different scales on the Y-axes in each row.

### Larval development time

Larval development time decreased with more larger workers in the low and high-low resource environments (Fig. 4d-e; LR test of smooth term vs constant: low – *χ*^*2*^ = 108.8, d.f. = 2.15, *P* < 0.001; high-low – *χ*^*2*^ = 200.7, d.f. = 2.71, *P* < 0.001). In the high resource environment, more workers of any size seemed to marginally increase development time (Fig. 4f; high - *χ*^*2*^ = 14.4, d.f. = 1.45, *P* < 0.001); however, these effects are negligible compared to the effects of worker size composition in the low and high-low resource environments (Fig. 4d-f).

### Larval survival

Larval survival decreased with more smaller workers in the low and high-low resource environments (Fig. 4g-h; low - *χ*^*2*^ = 78, d.f. = 2.2, *P <* 0.001; high-low - *χ*^*2*^ = 21.6, d.f. = 1.8, *P <* 0.001), but more of the largest observed workers had no effect on larval survival. Larval survival slightly decreased with more workers in the high resource environment, particularly with more larger workers (Fig. 4i; high – *χ*^*2*^ = 22.3, d.f. = 1.5, *P <* 0.001).

### Callow size

In the low resource environment, mean callow size increased with more larger workers and decreased with more smaller workers (Fig. 4j; *F* = 2.5, d.f. = 1.78, *P* = 0.018), but worker size composition was unrelated to standard deviation of callow size (Fig. 4m; low - *F* = 2.3E-7, d.f. = 3.9E-6, *P* = 0.74). In the high-low resource environment, worker size composition had little to no effect on mean callow size (Fig. 4k; *F* = 0.7, d.f. = 0.61, *P* = 0.068), whereas more larger workers slightly decreased the standard deviation in callow size (Fig. 4n; high-low - *F* = 2.6, d.f. = 1.75, *P* = 0.016). In the high resource environment, more workers of any size decreased the mean callow size (Fig. 4l; *F* = 17.3, d.f. = 1.7, *P* < 0.001), whereas worker size composition did not affect standard deviation in callow size (Fig. 4o; high - *F* = 0.7, d.f. = 0.66, *P* = 0.06).

## DISCUSSION

Size polymorphism in bumblebee colonies had different effects on worker production vital rates under different resource environments. Overall, colonies with more larger workers often had greater worker production compared to colonies with smaller workers. This pattern is similar to many performance-based (Goulson *et al*. 2002; Spaethe & Weidenmüller 2002; Peat & Goulson 2005; Ings 2007; Kapustjanskij *et al*. 2007; Spaethe *et al*. 2007) and manipulative experiments (Cnaani & Hefetz 1994). We found the opposite effect in two cases: more larger workers slightly decreased larval survival and more workers of any size decreased mean callow size in the high resource environment. We also never detected stronger per capita benefits of smaller workers relative to larger workers. We discuss each result in turn below, as well as some advantages and limitations of functional linear models.

### Functional implications of worker size distribution

Across social organisms, offspring number often increases with the number of helpers (Brown *et al*. 1982; Malcolm & Marten 1982; Biedermann & Taborsky 2011; Young *et al*. 2015), particularly when resources are high (Harrington, Mech & Fritts 1983; Doolan & Macdonald 1997). We found a similar impact on colony egg production in our high-low and high resource treatments, yet FLMs also revealed that in these environments more larger workers increased colony egg production relative to more smaller workers. Laboratory studies of bumblebees have shown that colonies consisting of only larger workers produce more eggs than colonies consisting of only smaller workers (Cnaani & Hefetz 1994). Larger workers are known to return more resources to the colony (Goulson *et al*. 2002; Kerr, Crone & Williams 2019), but they are less resilient against starvation (Couvillon & Dornhaus 2010). This tradeoff might explain why larger workers increased colony egg production only in the high-low and high resource environment. The opposite effect has been found in a fire ant, *Solenopsis invicta*, where monomorphic colonies of large workers produced almost no brood compared to monomorphic colonies of small workers (Porter & Tschinkel 1985). However, the size-based roles of workers in these two eusocial insects differs. Larger bumblebees are foragers (Cumber 1949; Goulson *et al*. 2002; Goulson 2009), but smaller fire ant workers do most of the foraging and feeding (Wilson 1978; Cassill & Tschinkel 1999). Larger fire ant workers live longer than smaller workers (Porter & Tschinkel 1985; Calabi & Porter 1989), which is the opposite of bumblebee workers (da Silva-Matos & Garofalo 2000; Kerr, Crone & Williams 2019). Therefore, the general mechanism may be similar, in spite of contrasting patterns.

In contrast to conclusions for egg production, more larger workers had either beneficial or neutral effects on development time and larval survival only in low and high-low environments, whereas more smaller workers had negative impacts on these vital rates. In bumblebees, there seems to be a resource-driven trade-off between provisioning for developing larvae and production of new eggs when resources are low; i.e., more larger workers under low resources maintain similar larval development time that is comparable to colonies under high resources at the cost of no apparent increase in egg production. Results for small workers in these colonies are similar to those for cooperative breeding species, in which the presence of more helpers often reduces offspring survival when resources are low (Harrington, Mech & Fritts 1983; Woodroffe & Macdonald 2000). These negative impacts of helpers in cooperative breeding species may be due to them shifting efforts towards increasing their own survival (Bruintjes, Hekman & Taborsky 2010), which seems less likely in bumblebees because workers are non-reproductive. Indeed, bumblebee workers are reported to switch from nursing to foraging tasks when resources are low (Cartar 1992), indicating that workers overall increase (not decrease) cooperative efforts. Additionally, bumblebee workers predominantly feed on nectar and larvae predominantly feed on pollen (Plowright & Pendrel 1977; Goulson 2009), which may reduce competition among siblings and enhance cooperative behaviors. It would be interesting to monitor foraging behavior of bumblebee workers during resource dearths, i.e. changes in nectar vs. pollen collection rates, to better understand their cooperative efforts. Life history and level of sociality might play a role in determining the cooperative efforts of individuals, where long lived species can afford to either reduce offspring number or cooperative behaviors towards offspring when resource are low.

Across our three environments, observed average size of all workers decreased in colonies with less available resources. In the low resource environment, more smaller workers resulted in callow workers of smaller sizes. Bumblebee workers have been recorded to be smaller on average in simple, intensively managed landscapes (Persson & Smith 2011). Laboratory experiments also show that colonies produce smaller workers during food shortages (Schmid-Hempel & Schmid-Hempel 1998). The correlation between worker size distribution and callow worker size suggests that stressful resource conditions may produce a negative feedback loop, where colonies of smaller workers cannot properly feed and care for brood (Cartar & Dill 1991) causing the emergence of smaller callow workers. Therefore, the cost and benefits of helpers within social groups may often regulate the traits of individuals (e.g. sex ratios, worker sizes) that are expressed (Griffin, Sheldon & West 2005). Functional linear models are only a correlative technique, so an alternative shared driver could be shifting the size distribution towards smaller workers. For example, lower resources could cause differential mortality of larger workers due to starvation (Couvillon & Dornhaus 2010) and cause larvae to develop into smaller callow workers because of fewer resources brought back by the remaining workers. Laboratory monomorphic colonies consisting of only small or large workers had no difference in the mean and variance in callow size when supplied with abundant resources (Cnaani & Hefetz 1994). If these laboratory colonies had to forage for resources and still produced workers of similar sizes, then we might be able to determine whether a shared driver is most likely causing these effects in our study.

### Functional linear models as a statistical approach in ecology

Previously, FLMs have been used to evaluate the lagged effects of environmental drivers on plant population dynamics (Teller *et al*. 2016; Tenhumberg *et al*. 2018). Here, we extend the use of FLMs to evaluate the size-based contribution of workers in bumblebee colonies. FLMs could be applied to understanding individual contributions in many high-dimensional social systems. For example, they could be used to explore the effects of age within social groups of different taxa and levels of sociality, such as eusocial honey bees (Seeley & Kolmes 1991), semi-social mole rates (Jarvis 1981; Zöttl *et al*. 2016), and cooperative breeding meerkats (Clutton-Brock *et al*. 2001a) or cichlid fish (Bruintjes & Taborsky 2011). In the African mole rat, larger groups had higher rates of offspring recruitment (Young *et al*. 2015) and cooperative behaviors were found to increase with age (Zöttl *et al*. 2016). Therefore, FLMs might be able to determine how vital rates (e.g. offspring recruitment) differ with the number of helpers of different ages for the African mole rate. FLMs provide an alternative way to study these high-dimensional ecological systems using field observational data, particularly where manipulative experiments may not be possible.

Correlative techniques, such FLMs, provide a valuable complement to many manipulative experiments that aim to test similar hypotheses. However, these separate approaches have their own set of advantages and limitations that need to be considered when making conclusions about these high-dimensional systems such as lagged effects or size polymorphism. For example, FLMs can be data-heavy (e.g., 20-25 independent observations of the signal and response; Teller *et al*. 2016); only inform us about correlations and not causations; and may have collinear predictors that obscure the true driver of these responses.

Collinearity is not specific to FLMs but is equally problematic for many simple (e.g. multiple regression) and complex statistical techniques (e.g. structural equation models). To date, only two studies have reported applying functional smoothing approaches to high-dimensional ecological systems by exploring how lagged environmental drivers influence plant performance (Teller *et al*. 2016; Tenhumberg *et al*. 2018). Teller et al. (2016) predicted how lagged effects of past precipitation and local competition influenced plant growth and survival; however, they would not be able to parse out the true driver of plant performance if density and precipitation covaried across some gradient. In our study, colony age and worker size composition had similar effects for half of our vital rates (n = 8 of 15) across the three resource environments. When exploring the trends and collinearity for these several vital rates (Appendix S4), two of four vital rates (Table 1) had confounding effects of colony age and size composition suggesting that either or both might be driving these trends (Table 1). When using simple or complex correlative methods, it is important to explicitly evaluate the collinearity of predictor variables as we have demonstrated here.

### Summary

Overall, we found that the advantages and disadvantages of workers of different sizes on worker production only became apparent when exploring these effects across these three different resource environments. We also found that bumblebee colonies shifted their worker size distribution across these resource environments. Among eusocial insects, caste size polymorphism is hypothesized to be an adaption to expand accessibility of resources, such as seed size in ants (Davidson 1978; Traniello & Beshers 1991; Retana & Cerdá 1994) and flower size in bumblebees (Peat, Tucker & Goulson 2005). However, the shift in worker size distribution across these resource environments could have emerged from the lower tolerance of larger workers to starvation (Couvillon & Dornhaus 2010). Prior to this study, quantifying the contribution of individuals in social groups has been challenging. Here, we demonstrate that functional linear models are not constrained to evaluating lagged effects on individual performance, but these models are a valuable complement to manipulative experiments and have the potential to explain many complex, trait-based life histories of social organisms.

## ACKNOWLEDGEMENTS

Our study was jointly funded by Tufts Graduate Student Research Award and Collaborative NSF Grant to NM. Williams (DEB1354022) and E.E. Crone (DEB1411420). We thank Nick Dorian, Sonja Glasser, Colin Fagan, Jessica Drost, Andrew Buderi, and John Mola for field assistance, and the Laidlaw Bee Biology Facility at UC Davis for use of their facilities.

## AUTHOR CONTRIBUTIONS

NZK, EEC and NMW conceived the ideas and designed methodology; NZK, RLM, and NMW collected the data; NZK and EEC analyzed the data; and NZK, EEC, RLM, and NMW wrote the manuscript.

