## Supplementary Information for "Bumblebee worker body size affects new worker production in different resource environments"

**Figure S1.** Google earth aerial images of the 2015 and 2016 field sites at UC Davis.

**Figure S2.** Scatterplot matrix of colony age, mean and standard deviation in worker size, and population size of colonies in the low resource environment.

**Figure S3.** Scatterplot matrix of colony age, mean and standard deviation in worker size, and population size of colonies in the high-low resource environment.

**Figure S4.** Scatterplot matrix of colony age, mean and standard deviation in worker size, and population size of colonies in the high resource environment.

**Appendix S1.** Exploring effects of colony age on vital rates using GAMs.

**Appendix S2.** Exploring the effects of worker size composition on vital rates when standardized by worker production costs.

**Appendix S3.** Calculations of model predicted values for generalized additive models.

**Appendix S4.** Evaluating confounding effects of colony age and worker size composition on vital rates affecting worker production

**Figure S1.** Google earth aerial images of the field sites at UC Davis during experiment periods using package ‘plotKML’ (Hengel et al. 2015) in program R. A larger landscape view (left) and smaller scale view (right) of each of the study sites. The 2015 colonies (a-b) were located at Harry H. Laidlaw Honey Bee Research (red box) at UC Davis, where the surrounding landscape was agricultural land, floral research plots (orange boxes), and 0.2 ha bee-friendly garden (blue box). The 2016 colonies were located (c-d) in agricultural fields on UC Davis Experimental Farm property.

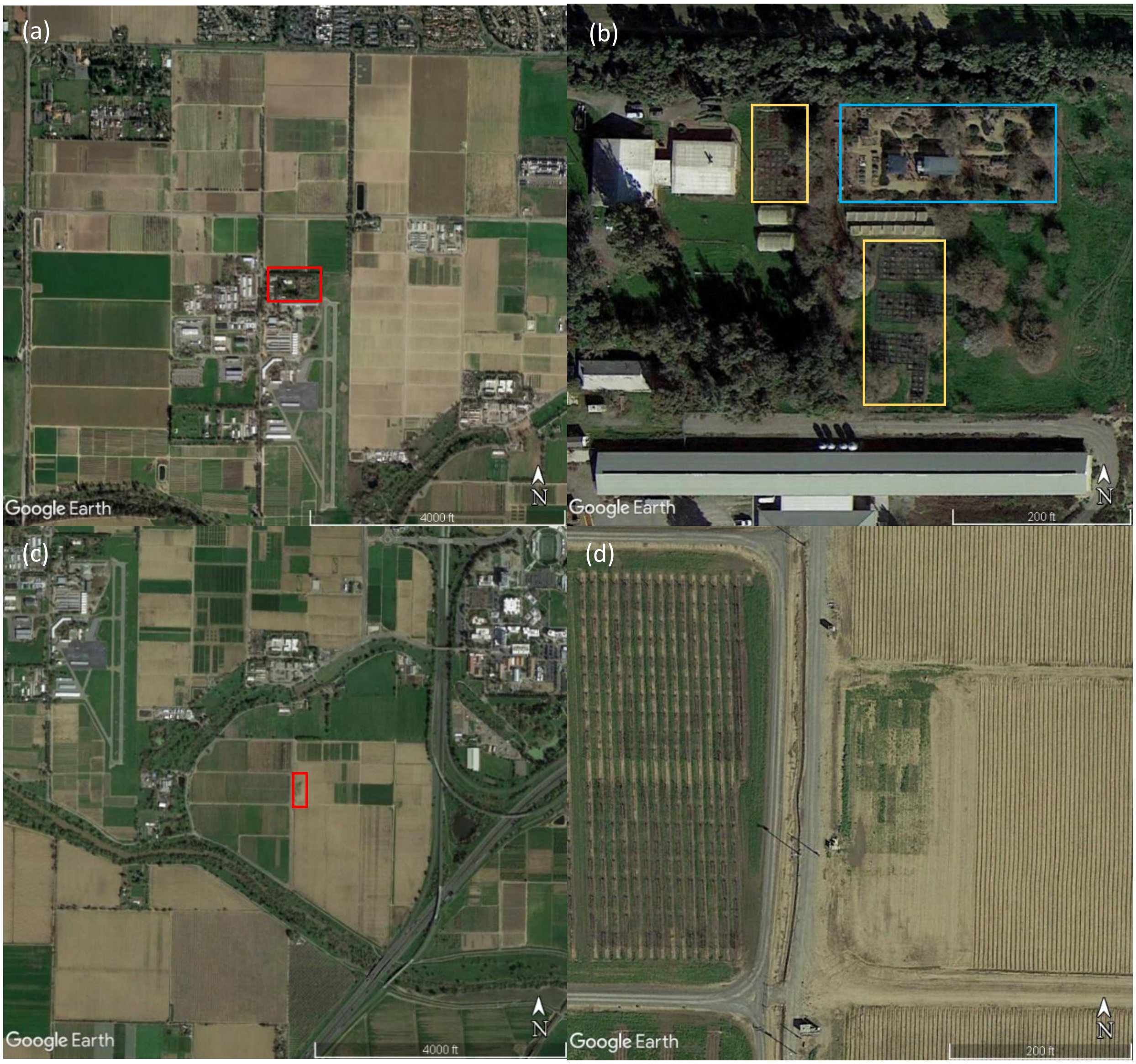

**Figure S2.** Scatterplot matrix of mean and standard deviation (sd) in worker size measured as intertegular span in mm (ITS), colony age, and population size of colonies in the low resource environment. The leading diagonal contains a histogram a of each variable, the upper diagonal panels contains the correlation coefficient between each pairwise comparison, and the lower diagonal panel contains a scatterplot with the mean and 95% confidence intervals between each pair.

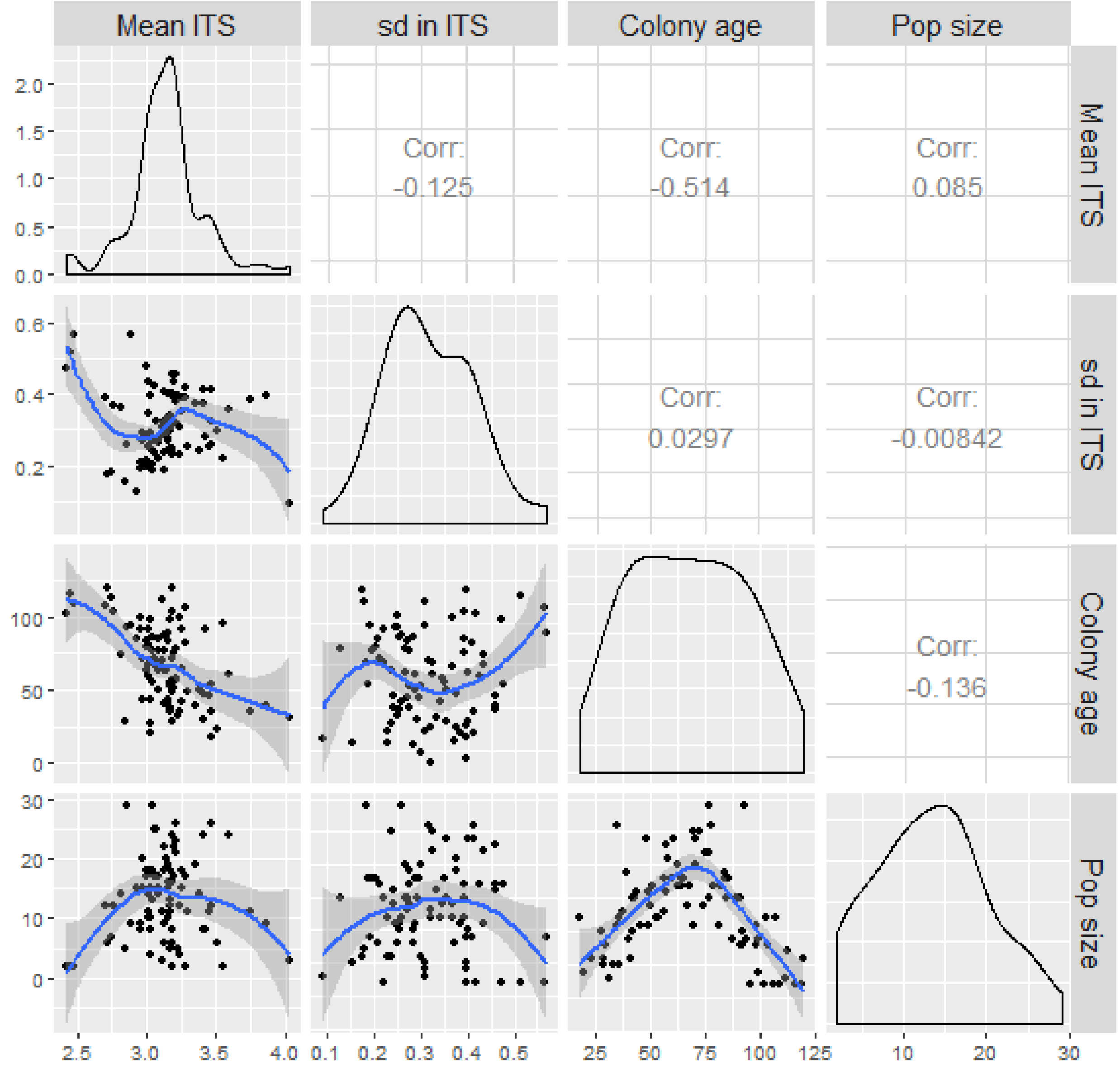

**Figure S3.** Scatterplot matrix of mean and standard deviation (sd) in worker size measured as intertegular span in mm (ITS), colony age, and population size of colonies in the high-low resource environment. The leading diagonal contains a histogram a of each variable, the upper diagonal panels contains the correlation coefficient between each pairwise comparison, and the lower diagonal panel contains a scatterplot with the mean and 95% confidence intervals between each pair.

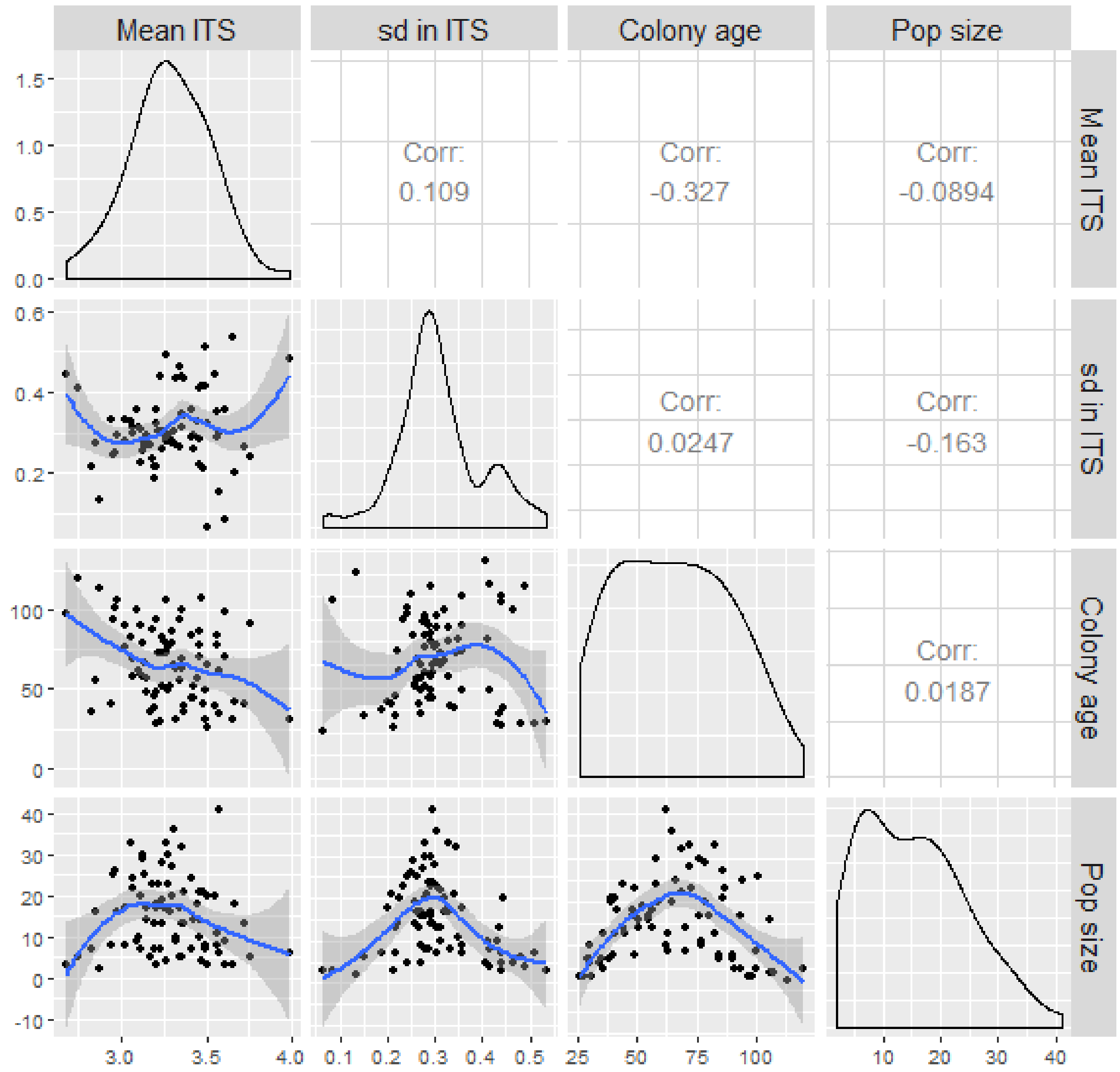

**Figure S4.** Scatterplot matrix of mean and standard deviation (sd) in worker size measured as intertegular span in mm (ITS), colony age, and population size of colonies in the high resource environment. The leading diagonal contains a histogram a of each variable, the upper diagonal panels contains the correlation coefficient between each pairwise comparison, and the lower diagonal panel contains a scatterplot with the mean and 95% confidence intervals between each pair.

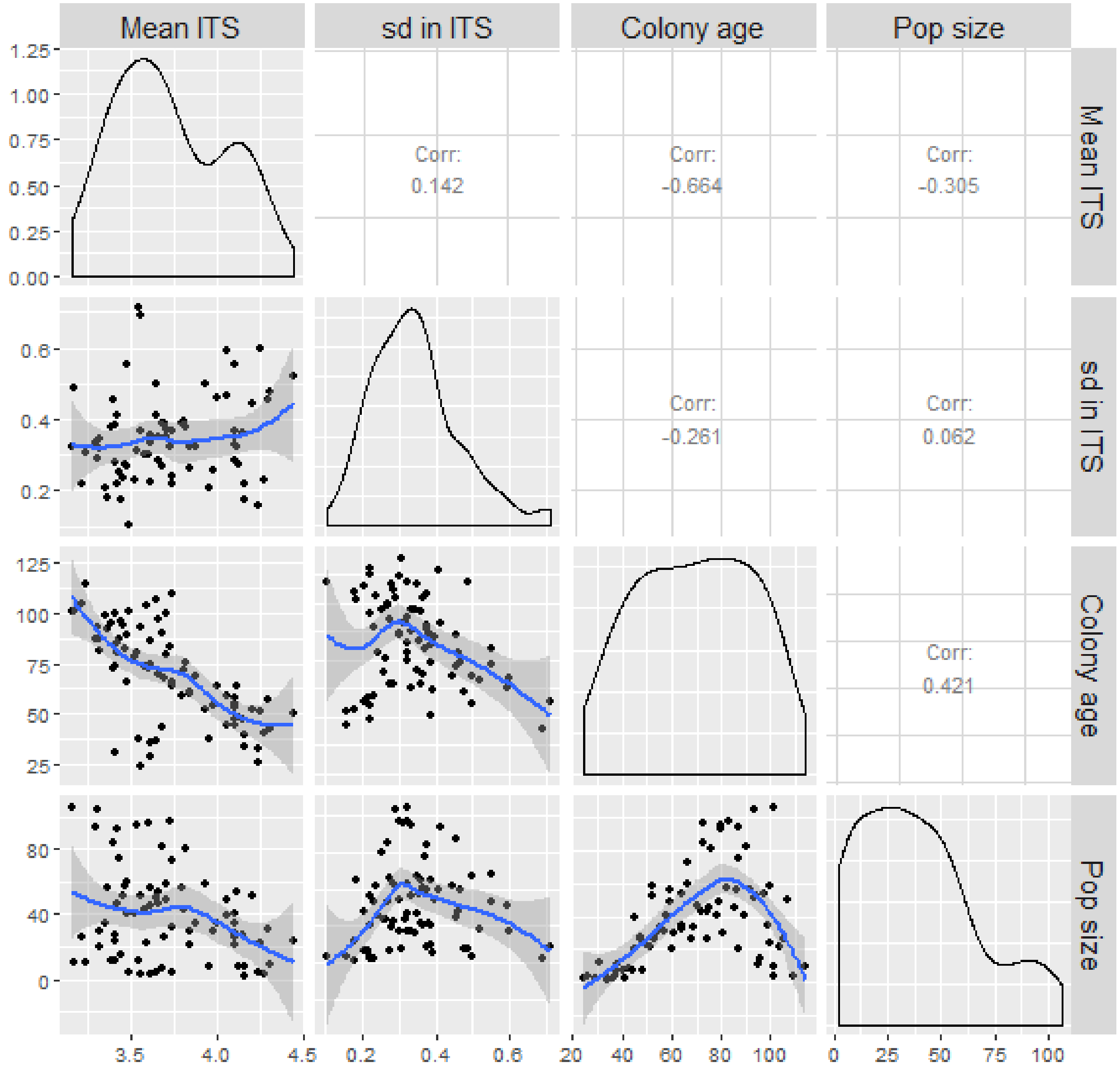

**Appendix S1.** Exploring effects of colony age on vital rates using GAMs.

Methods

In addition to the FLMs for worker size composition (see main text), we fit a second set of GAMs predicting each response as a smooth covariate of colony age. For new eggs laid, larval development time, and larval survival, we knew when eggs first appeared and used the colony age at this time of first detection. We estimated time of first detection for each callow worker in each colony, and we used the date of emergence minus average development time for each colony to estimate colony age. We used the same probability distributions as the worker size composition FLMs for evaluating colony age effects. We did not restrict the number of knots for colony age GAMs since lab and field conditions may alter the effect of colony age on vital rates.

Results

In the low resource environment, daily egg production remained constant with increasing colony age until 60 days old at which point egg production steadily declined (Fig. S1.1a; significance of colony age smooth term: $\chi^2$ = 13.4, d.f. = 1.76, $P$ < 0.001). In the high-low resource environment, egg production increased with colony age until a switch point around 60 days since the appearance of the first brood mass, where egg production rapidly declined (Fig. S1.1b; $\chi^2$ = 21.5, d.f. =1.9, $P$ < 0.001). Egg production remained constant with colony age in the high resource environment (Fig. S1.1c; $\chi^2$ = 1E-4, d.f. = 4.1E-4, $P$ = 0.65).

In the low resource environment, larval development time initially decreased with increasing colony age then increased after 60 days (Fig. 1.1d; $\chi^2$ = 202.8, d.f. = 3.96, $P$ < 0.001). In the high-low environment, larval development increased with colony age until approximately 60 days and then decreased (Fig. S1.1e; $\chi^2$ = 153.8, d.f. = 1.98, $P$ < 0.001). Larval development time remained constant with colony age in the high resource environment (Fig. S1.1f; $\chi^2$ = 3.3E-5, d.f. = 8.5E-7, $P$ = 0.55).

Larval survival decreased with colony age in both the low and high-low resource environments (Fig. S1.1g-h; low - $\chi^2$ = 83.1, d.f. = 1.7, $P$ < 0.001; high-low - $\chi^2$ = 166.4, d.f. = 1.8, $P$ < 0.001). In the high resource environment, larval survival remained constant with colony age (Fig S1.1i; $\chi^2$ = 15.6, d.f. = 1.6, $P$ < 0.001).

Mean callow size decreased with colony age in both low and high-low resource environments (Fig. S1.1j-k; low - $F$ = 3.2, d.f. = 1.05, $P$ = 0.009; high-low - $F$ = 5.9, d.f. = 2.56, $P$ < 0.001), whereas mean callow size remained approximately constant after colonies were placed in the high resource environment (Fig. S1.1l; $F$ = 22.7, d.f. = 3.15, $P$ < 0.001). Colony age had no effect on the standard deviation of callow size across resource environments (Fig. S1.1m-o; low - $F$ = 0.03, d.f. = 0.06, $P$ = 0.31; high-low - $F$ = 9.7E-6, d.f. = 4.7E-5, $P$ = 0.54; high - $F$ = 0.8, d.f. = 0.52, $P$ = 0.17).

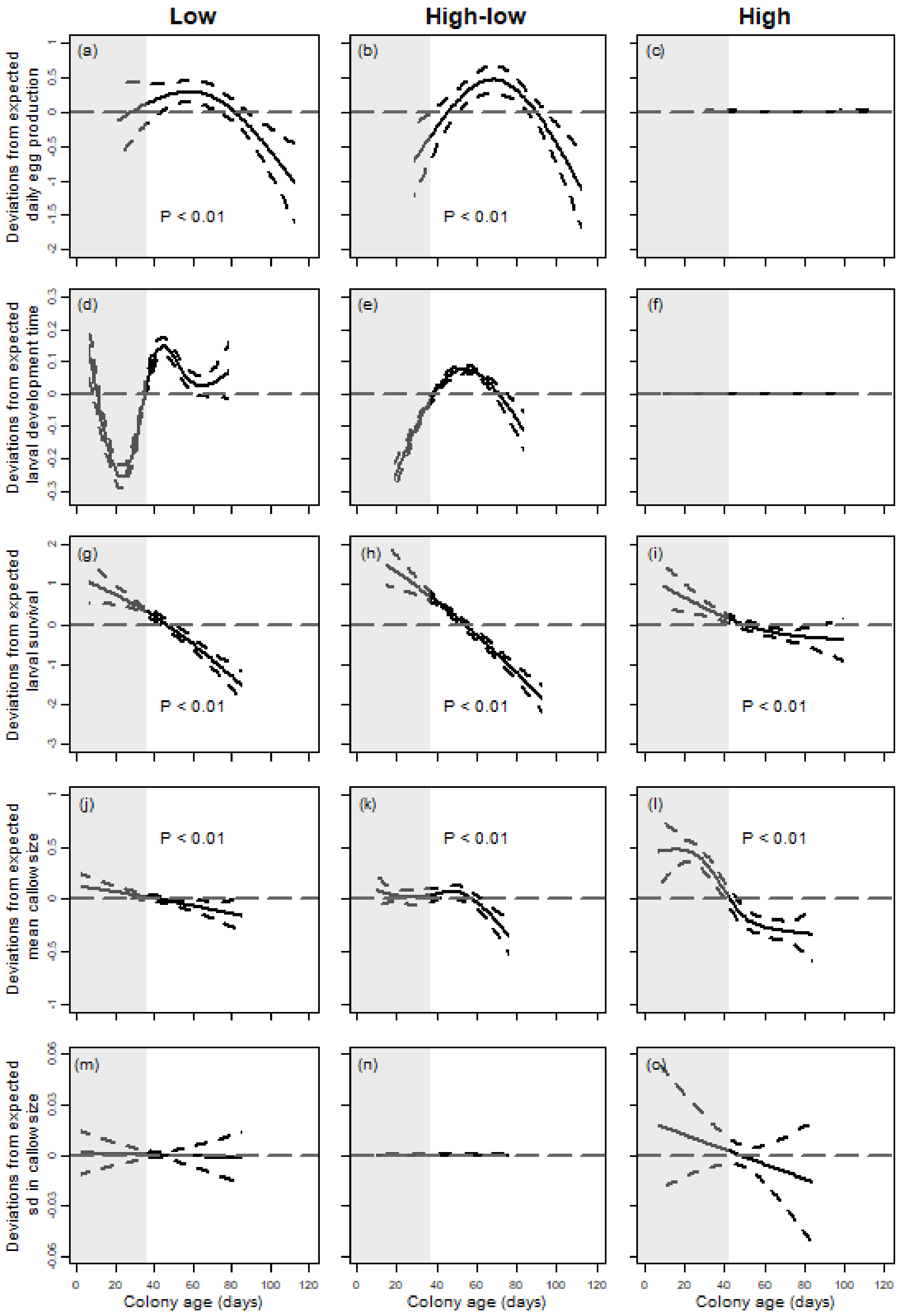


**Figure S1.1.** Generalized additive model results depicting the smooth function of the deviations from expected values for all five vital rates as a function of colony age in days for the low (left), high-low (middle), and high (right) resource environments. Dotted horizontal line represent deviations from mean values, i.e. values above the line have positive impact on Y and values below the line have negative effects. Lighter shaded and unshaded areas in the plot represent when the colonies were in the laboratory and outside in the field, respectively. Because all colonies experienced similar conditions in the lab, we focus on results during the time when they were in field environments (non-shaded area). Plots with a significant smooth term of colony age are labeled with $P < 0.01$. Note different scales on the Y-axes in each row.

**Appendix S2.** Exploring the effects of worker size composition on vital rates when standardized by worker production costs.

### Methods

Even if larger workers contribute more to the production of new workers on a per capita basis, smaller workers may have more benefits after accounting for differences in their production costs (c.f. Kerr, Crone & Williams 2019). Therefore, in addition to presenting FLMs as fitted (see Fig. 3), we also adjusted our worker size composition functions to reflect a constant mass of workers of each size (see Fig. 4). For example, if we found that more larger workers increased egg production, colonies producing only larger workers would pay a cost of having fewer workers due to their higher respective production costs. To adjust these functions by size-based production costs of workers, we converted slopes (change in response per worker) to change in response per mass. Specifically, we divided size-specific slopes, $\beta(s_x)$, by mass per worker in each size class, using the mass function reported by Kerr et al. (2019). Here, we present FLM results that account for worker production costs, to evaluate whether the size-based cost of worker biomass would affect contributions of different-sized workers.

### Results

#### *Egg production*

When accounting for production costs of larger workers, more larger workers still increased egg production in both resource environments. For workers > 4 mm, the effect of worker size on egg production was negligible (Fig. S2.1b-c).

#### *Larval development time*

Larval development time increased even more with more smaller workers in the low and high-low resource environments when accounting for production costs (Fig. S2.1d-e). Larvae took longer to develop with more intermediate-sized workers compared to more larger workers in the low resource environment (Fig. S2.1d), but larval development time was similar with either intermediate-sized or larger workers in the high-low environment (Fig. S2.1e). More small workers increased development time in the high resource environment when accounting for production costs (Fig. S2.1f), but the magnitude of this effect was negligible for observed worker sizes (see Fig. S3.3f).

#### *Larval survival*

Larval survival increased with more larger workers, even after accounting for production costs (Fig. S2.1g-h). In the high resource environment, more workers decreased larval survival, particularly with more larger workers, even when accounting for production costs (Fig. S2.1i). However, the magnitude of this effect was small over the realized range of worker size distributions (Fig. S3.3i).

*Callow size*

More larger workers increased the mean callow size in the low resource environment (Fig. 2); an effect that was exaggerated when we accounted for production cost (Fig. S2.1j). Worker size composition had no effect on the standard deviation in callow size in the low resource environment, even when accounting for production costs (Fig. S2.1m).

Worker size composition had little to no effect on mean callow size in the high-low resource environment (Fig. 4k), but more larger workers slightly decreased mean callow size when we accounted for production costs (Fig. S2.1k). More larger workers slightly decreased the standard deviation in callow size in high-low resource environment (Fig. 4n); an effect that was exaggerated when accounting for production costs (Fig. S2.1n).

When accounting for production cost, more workers of any size still decreased the mean callow size in the high resource environment (Fig. S2.1). No effects of worker size composition on standard deviation in callow size were seen in the high resource environment (Fig. 2o), yet standard deviation in callow size in the high resource environment slightly decreased with more workers when accounting for production costs, particularly more larger workers (Fig. S2.1o).

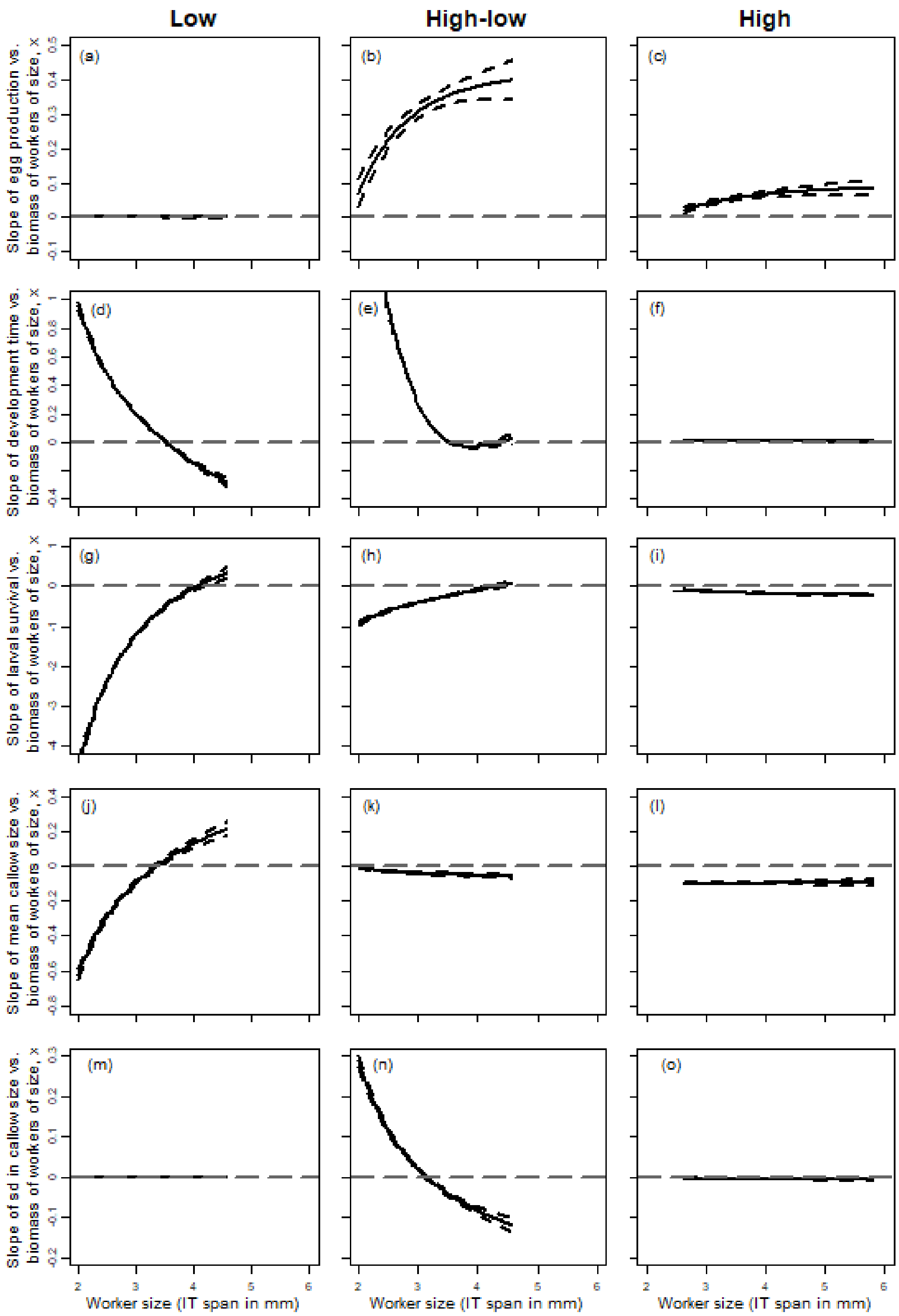


**Figure S2.1.** Generalized additive model results standardized by worker production costs (i.e. worker biomass) depicting the smooth function of the slopes for all five vital rates versus the number of workers of size x per worker biomass as a function of worker size x for the low (left), high-low (middle), and high (right) resource environments. Worker size was measured as the distance between the tegula, i.e. intertegular (IT) span in mm. Grey dotted horizontal line at zero represent deviations from mean slope values per worker biomass, i.e. slopes above the line means more workers of size x have positive impact on Y. Note different scales on the Y-axes in each row.

**Appendix S3.** Calculations of model predicted values for generalized additive models.

Methods

To calculate model predicted values for the five vital rates from the functional linear models (Fig. S2.1), we wanted to adjust colony size to account for size-based production costs of workers (i.e. colonies might have more workers if they produced only small workers or less workers if they produced only large workers). To do so, we first assumed that the total mass of the observed population sizes reflects the mass of the average worker size. Under this assumption, we calculated the mass of each observed population size (population mass of average sized workers in grams = population size × mass of average worker size) for colonies in all three resource environments. We calculated the mass of the average-sized worker (IT span; low – 3.16 mm; high-low – 3.31 mm; high – 3.68 mm) across resource environments, given relationship between IT span and weight [Intercept: - 0.036, slope: 0.044] (Kerr, Crone, & Williams 2019). Then, we recalculated the predicted population size of each worker size class in our FLMs (that was defined by IT spans) based off their respective worker mass (i.e. predicted population size of workers of size $x$ = population mass of average-sized workers / mass of worker size $x$). We then used these predicted population sizes for each worker size to calculate model predicted values (e.g. daily egg production) of hypothetical colonies consisting only of workers of each size class (see Fig. S3.3).

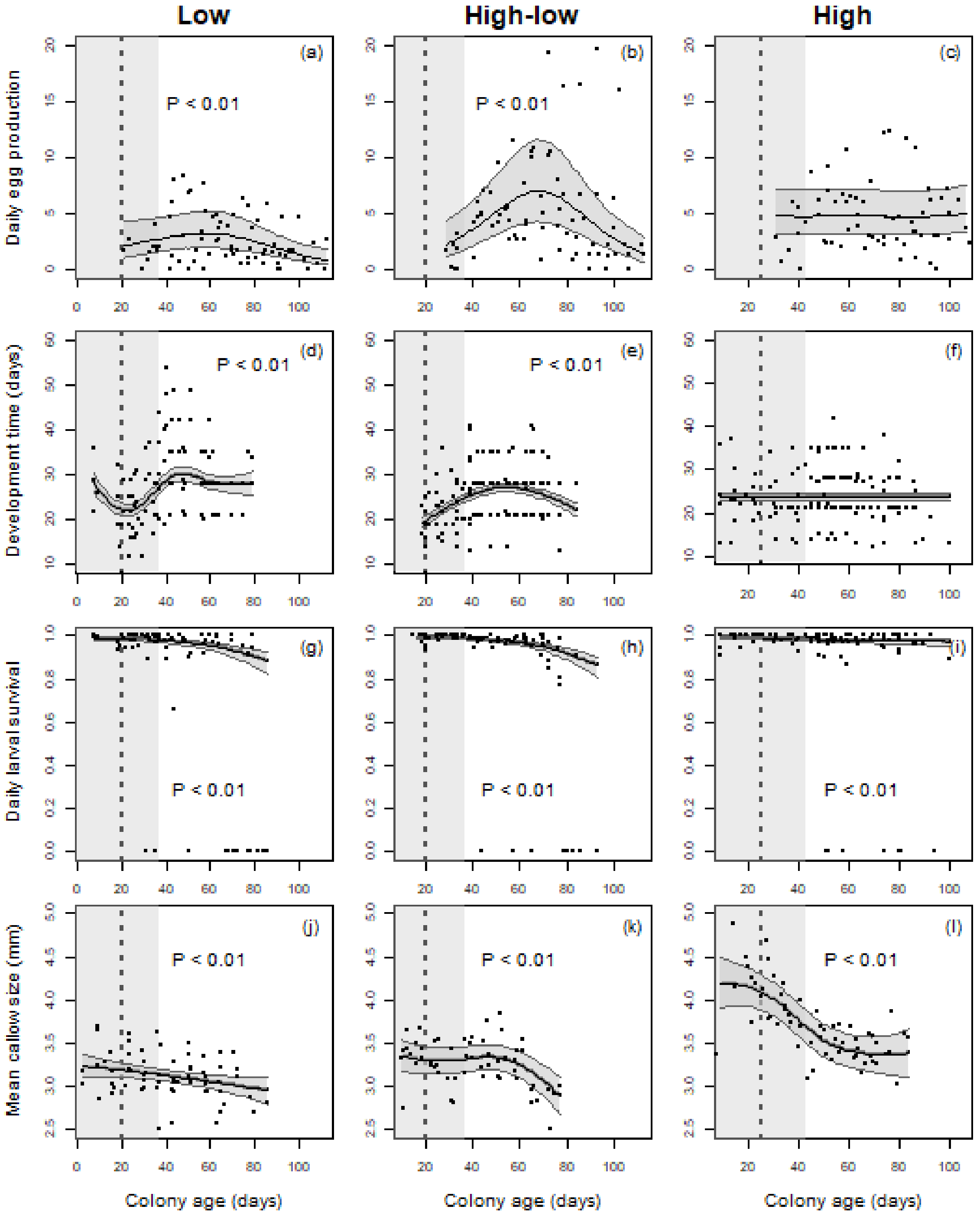


**Figure S3.1.** The effects of colony age of daily egg production (a-c), larval development time in days (d-f), daily larval survival (g-i), and mean callow size, which is measured as intertegular span in mm (j-l) in the low, high-low, and high resource environments. Darker shaded polygon around the mean functions represent the 95% confidence intervals. The vertical dotted line represents the appearance of the first worker cohort. Lighter shaded and unshaded areas in the plot represent when the colonies were in the laboratory and outside in the field, respectively. Because all colonies experienced similar conditions in the lab, we focus on results during the time when they were in field environments (non-shaded area).

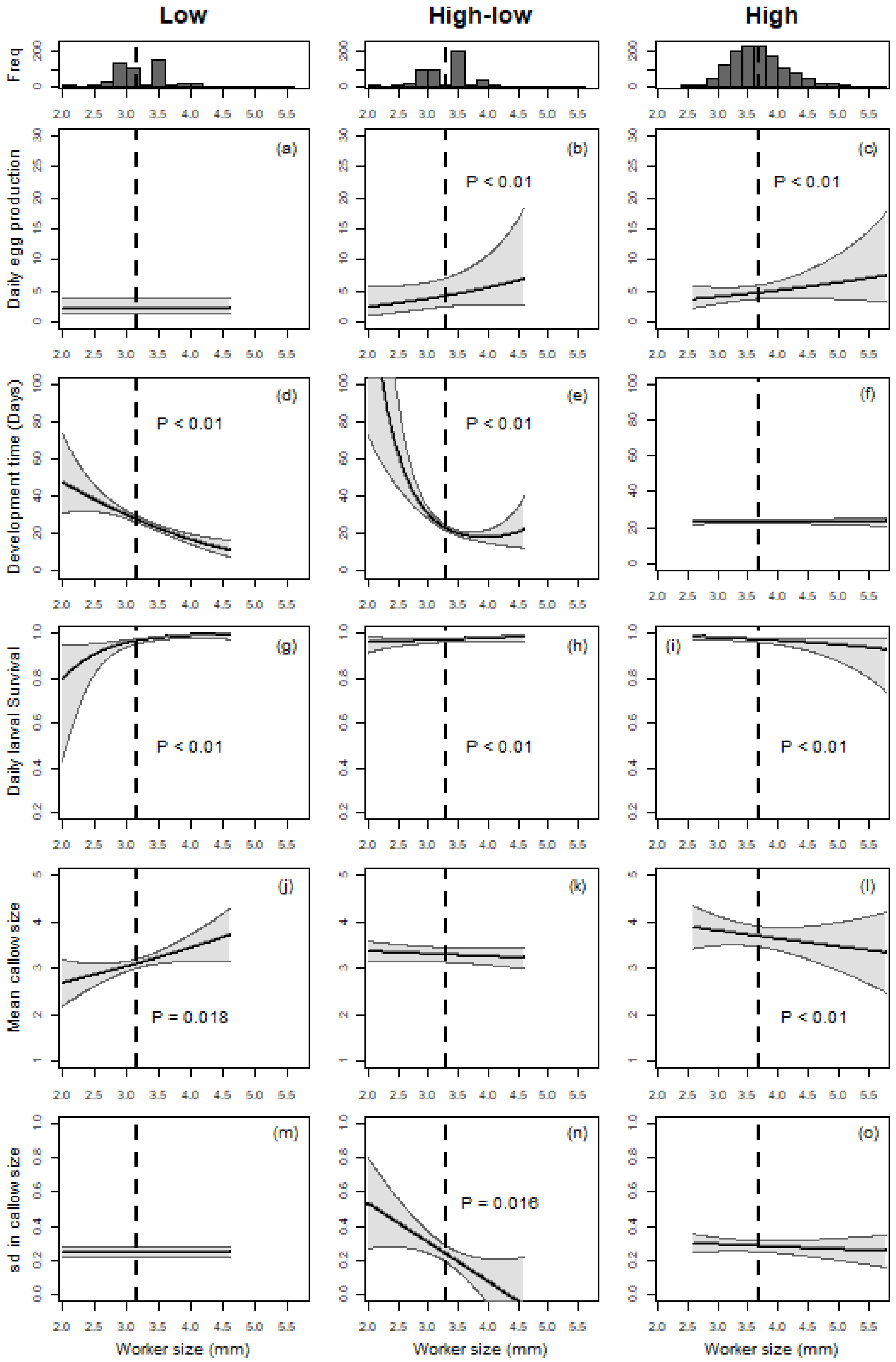


**Figure S3.2.** The contribution of workers of size *x* to daily egg production (a-c), larval development in days (d-f), daily larval survival (g-i), and mean callow size (j-l) in the three resource environments. The shaded areas represent the 95% confidence intervals around the mean. Histograms represent the worker size distribution across the season for all colonies. The vertical dotted line represents the average worker size across colonies in each treatment.

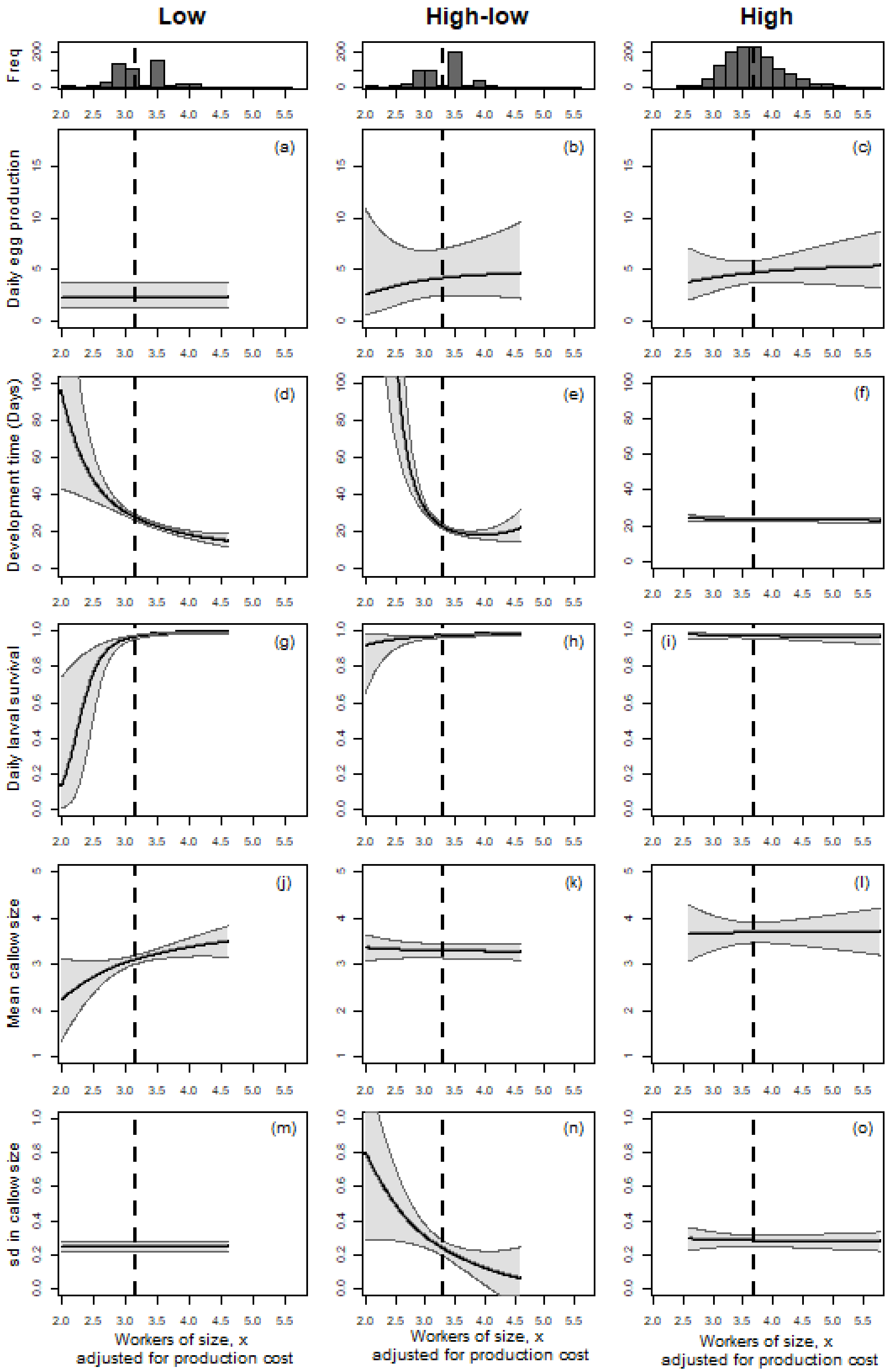

**Figure S3.3.** The contribution of workers of size *x* standardized by their production costs to daily egg production (a-c), larval development in days (d-f), daily larval survival (g-i), and the mean (j-l) and standard deviation (m-o) in callow size across the three resource environments. We offset the population size of workers of size *x* by their respective mass to account for their production costs, assuming cost of producing workers is proportional to their mass. The shaded areas represent the 95% confidence intervals around the mean. Histograms represent the worker size distribution across the season for all colonies. The vertical dotted line represents the average worker size across colonies in each treatment.

**Appendix S4.** Evaluating confounding effects of colony age and worker size composition smooth terms on five vital rates affecting worker production.

1. *Egg production*

Low: Colony age had no significant effects on egg production in the low resource environment. Therefore, we did not need to evaluate confounding effects for this vital rate.

High-low: Colony age and worker size composition both had significant effects on egg production in the high-low resource environment. When exploring confounding effects, we found that egg prodution is the highest at a colony age of ~ 60 days (Fig S4.1c), which is when colony size was the highest (Fig S4.1a). However, more larger workers contribute the most towards egg production (Fig S4.1d), which were found at younger colony ages (Fig S4.1b). Regardless, more workers of most worker sizes increases egg production (Fig S4.1d) suggesting that these smooth terms might be confounded.

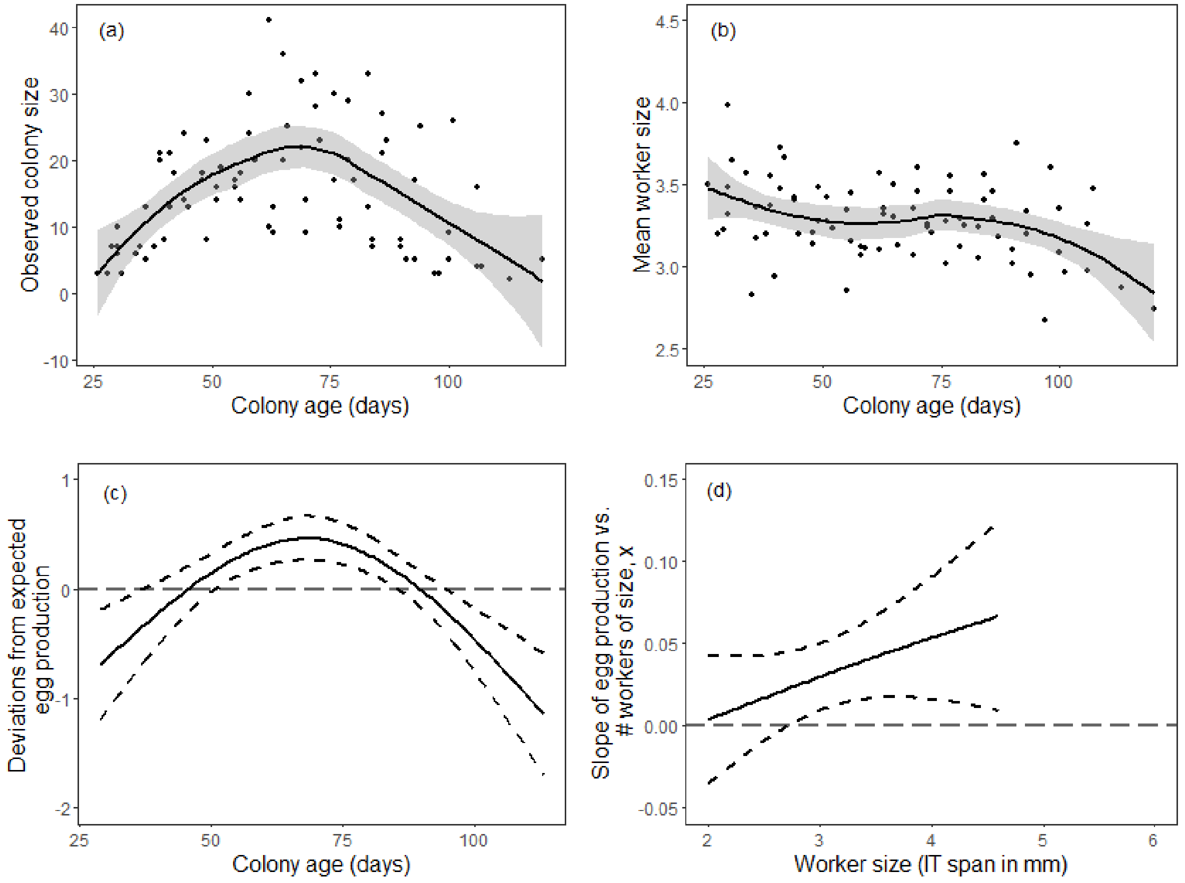


**Figure S4.1.** The (a) observed colony size (i.e. number of workers) and (b) mean worker size plotted against colony age in the high-low resource environment. Smooth components of generalized linear models evaluating the (c) deviations from expected egg production as a function of colony age and (d) the slope of egg production vs. # of workers as a function of worker size *x* for the high-low resource environment.

High: More larger workers increased egg production in the high resource environment but colony age had no effect. Therefore, we did not need to evaluate confounding effects for this vital rate.

2. *Development time*

Low: Colony age and worker size composition both had significant effects on egg production in the low resource environment. Development time increased with colony age and plateaued after 60 days (Fig S4.2c). More smaller workers also increased development time (Fig S4.3d), and worker size was the lowest at older colony ages (Fig S4.2b) where development time was the highest (Fig S4.2a). Therefore, colony age and WSC might have confounding effects. However, these colony age patterns on development time are likely attributed to lab conditions. Due to higher resource availability in the lab compared to the field, more larger workers eclosed and larvae developed faster. Since development time stabilized once in the field, we would contribute these early colony age patterns to worker size and feeding regiments in the lab.

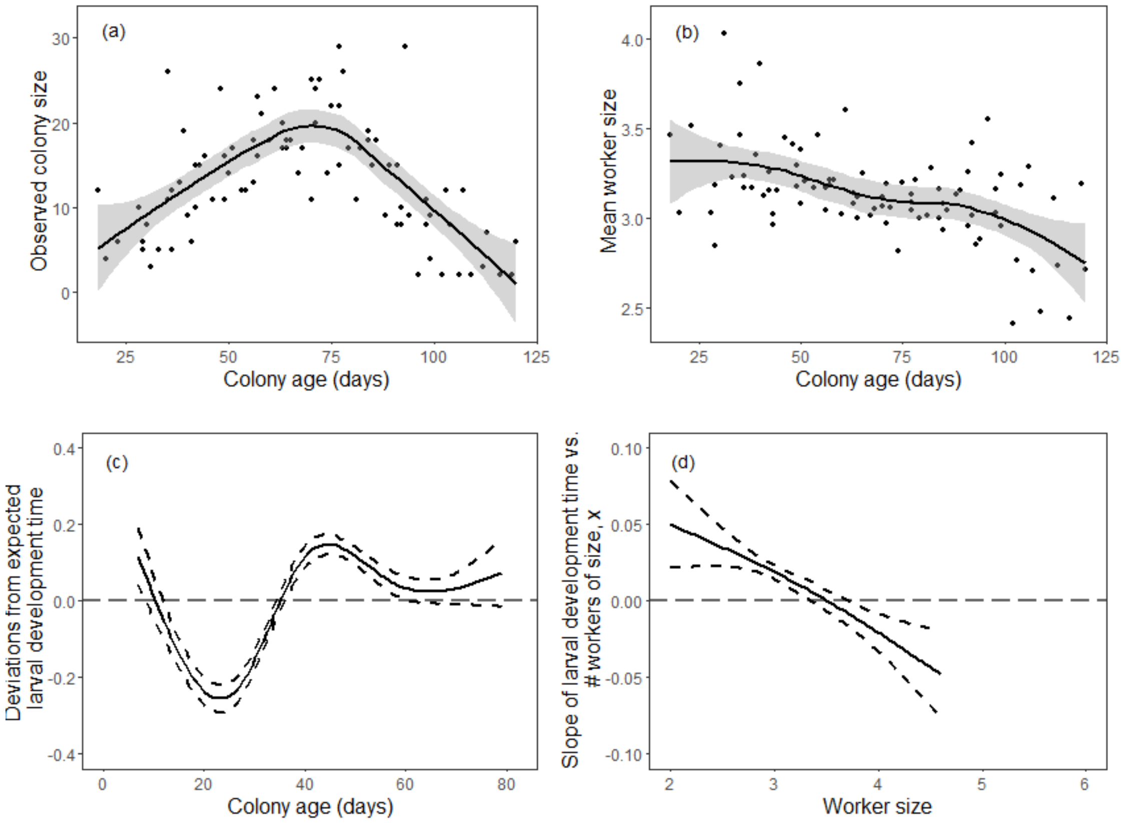


**Figure S4.2.** The (a) observed colony size (i.e. number of workers) and (b) mean worker size plotted against colony age in the low resource environment. Smooth components of generalized linear models evaluating the (c) deviations from expected egg production as a function of colony age and (d) the slope of egg production vs. # of workers as a function of worker size *x* for the low resource environment.

High-low: Colony age and worker size composition both had significant effects on egg production in the high-low resource environment. Development time was the highest around 50 days old (Fig S4.3c), which is the intermediate colony age. More smaller workers increased development time (Fig S4.3d), and worker size was the lowest at older colony ages (Fig S4.3b). Therefore, colony age and WSC might have confounding effects. However, similar to the low resource environment, the feeding conditions in the laboratory might have decreased development and increased worker size.

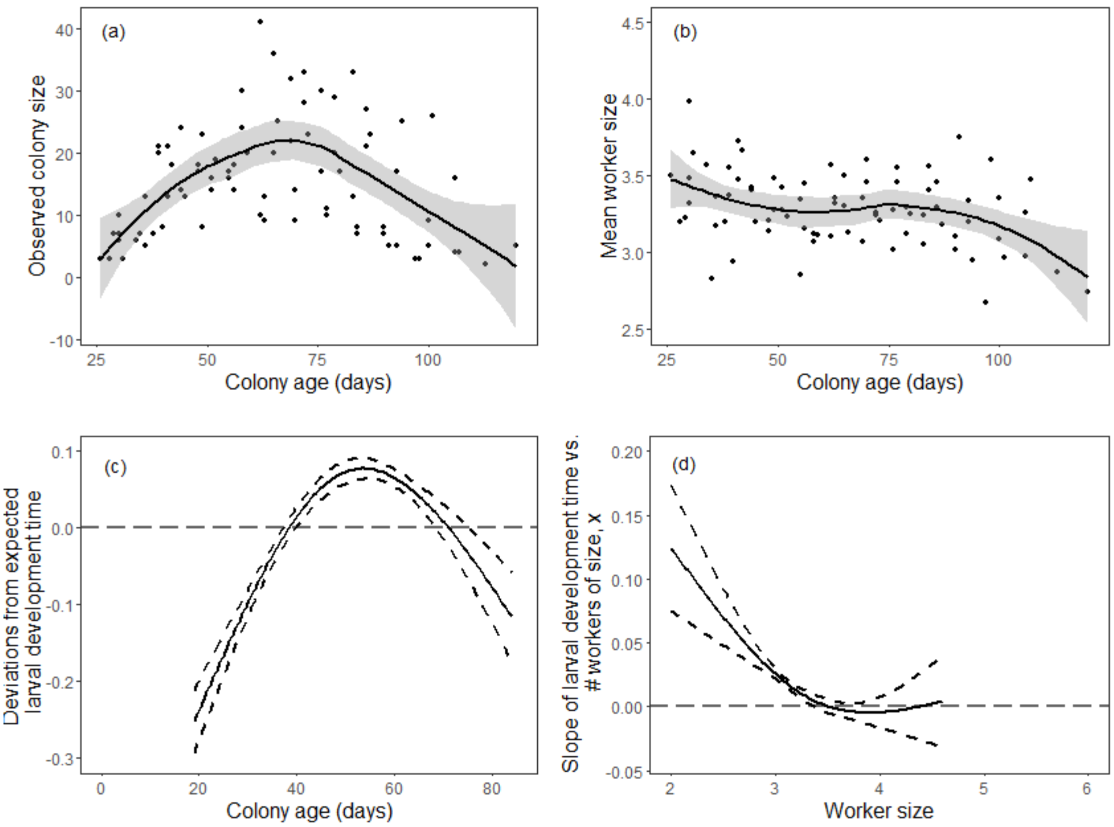


**Figure S4.3.** The (a) observed colony size (i.e. number of workers) and (b) mean worker size plotted against colony age in the high-low resource environment. Smooth components of generalized linear models evaluating the (c) deviations from expected larval development time as a function of colony age and (d) the slope of larval development time vs. # of workers as a function of worker size *x* for the high-low resource environment.

<u>High:</u> Neither colony age or worker size contribution had significant effects on development time in the high resource environment. Therefore, we did not need to evaluate confounding effects for this vital rate.

3. *Larval survival*

<u>Low:</u> Larval survival decreased with more smaller workers (Fig S4.4d) and at older colony ages (Fig S4.4d). Since worker size is lowest at older colony ages (Fig S4.4b), this suggests that colony age and WSC have confounding effects.

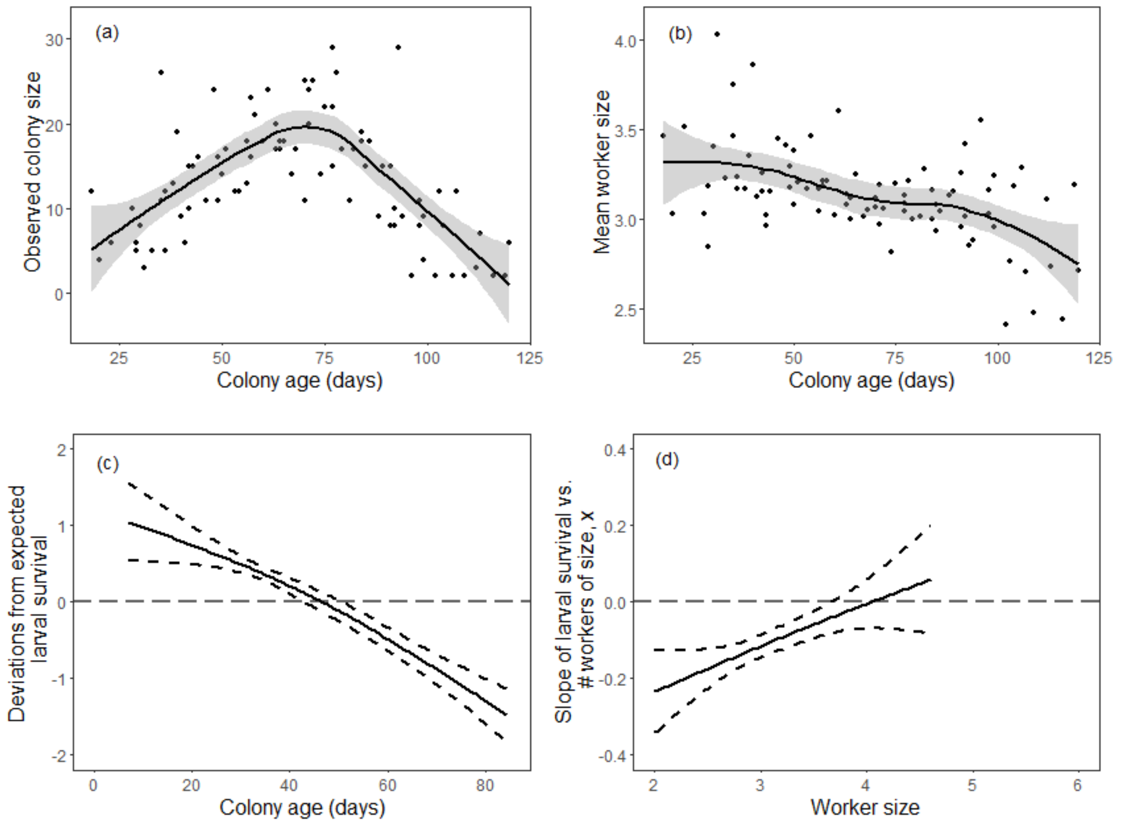


**Figure S4.4.** The (a) observed colony size (i.e. number of workers) and (b) mean worker size plotted against colony age in the low resource environment. Smooth components of generalized linear models evaluating the (c) deviations from expected larval survival as a function of colony age and (d) the slope of larval survival vs. # of workers as a function of worker size *x* for the low resource environment.

High-low: Larval survival decreased with more smaller workers (Fig S4.5d) and at older colony ages (Fig S4.5c). Since worker size was the lowest at older colony ages and highest at younger colony ages (Fig S4.5b), this suggests that colony age and WSC have confounding effects.

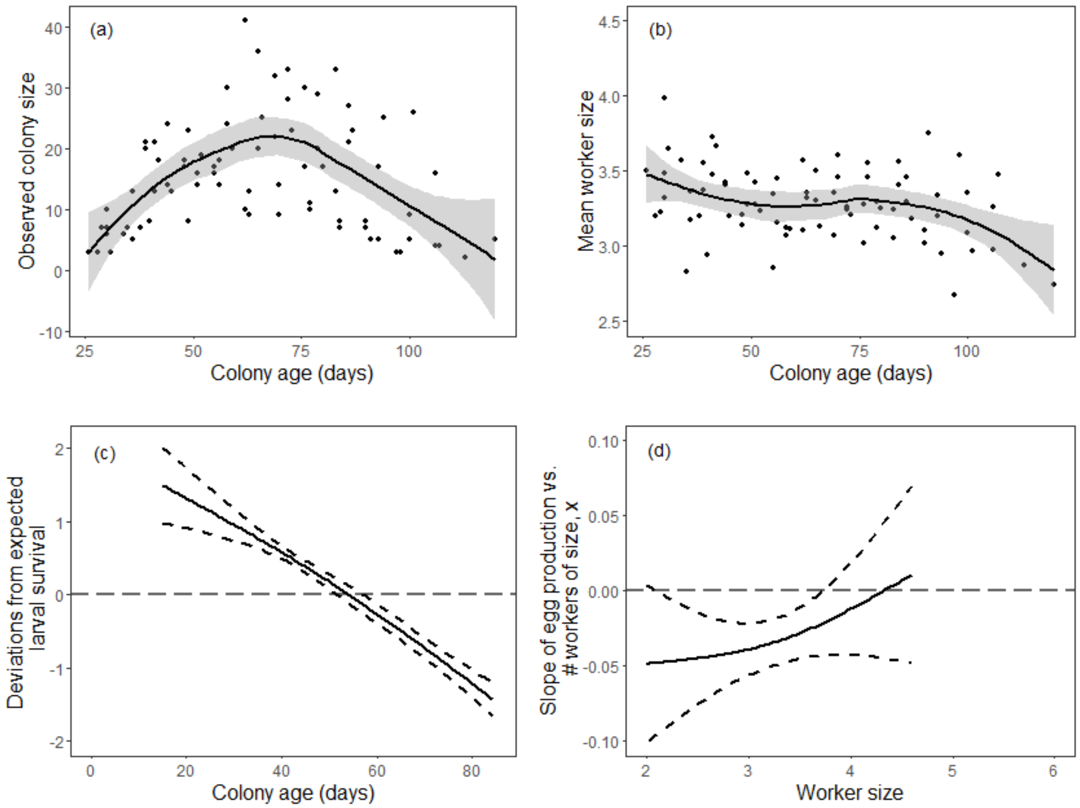


**Figure S4.5.** The (a) observed colony size (i.e. number of workers) and (b) mean worker size plotted against colony age in the high-low resource environment. Smooth components of generalized linear models evaluating the (c) deviations from expected larval survival as a function of colony age and (d) the slope of larval survival vs. # of workers as a function of worker size *x* for the high-low resource environment.

<u>High:</u> Larval survival decreased with increasing colony age (Fig S4.6c) and more larger workers (Fig S4.6d). Since worker size was the highest at younger colony ages (Fig S4.6b), this suggests that colony age and WSC do not have confounding effects.

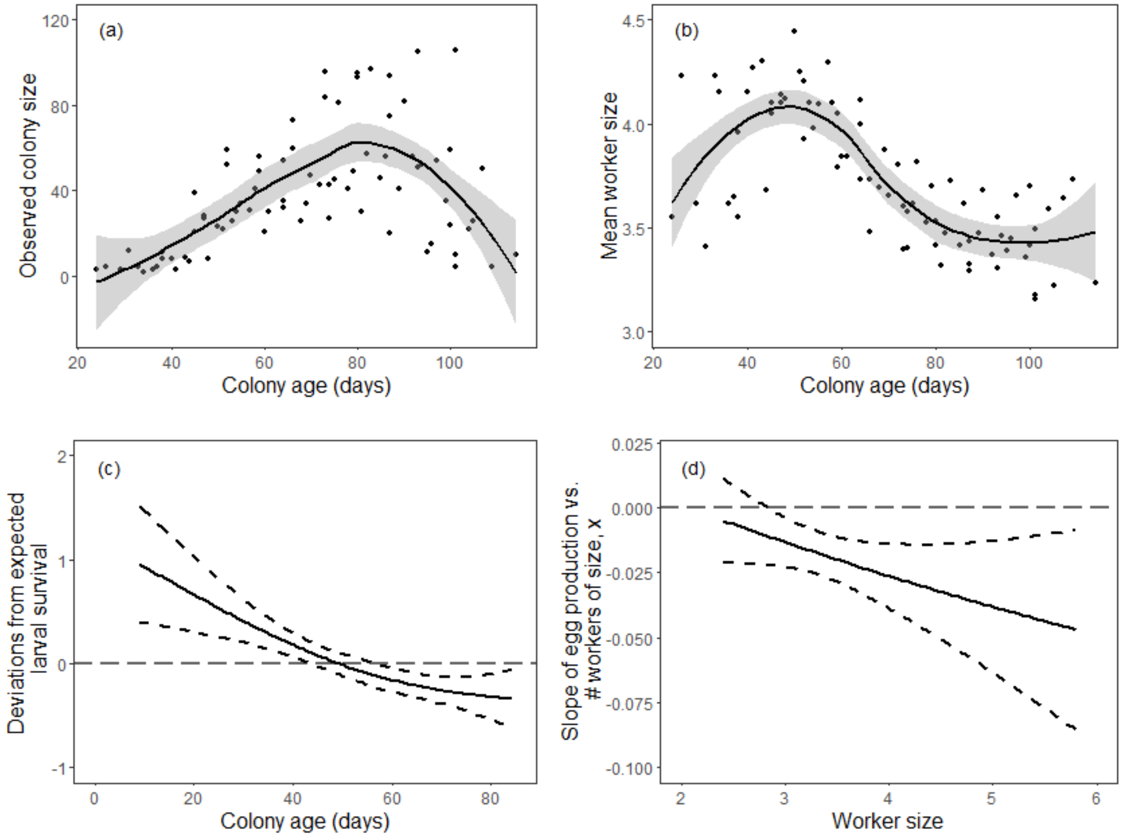


**Figure S4.6.** The (a) observed colony size (i.e. number of workers) and (b) mean worker size plotted against colony age in the high resource environment. Smooth components of generalized linear models evaluating the (c) deviations from expected larval survival probability as a function of colony age and (d) the slope of larval survival vs. # of workers as a function of worker size *x* for the high resource environment.

#### 4. *Mean callow size*

Low: Mean callow size decreased with increasing colony age (Fig S4.8c), but marginally increased with more larger workers (Fig S4.8d). Since worker size was the highest at younger colony ages (Fig S4.8b), these smooth terms are likely to be confounding.

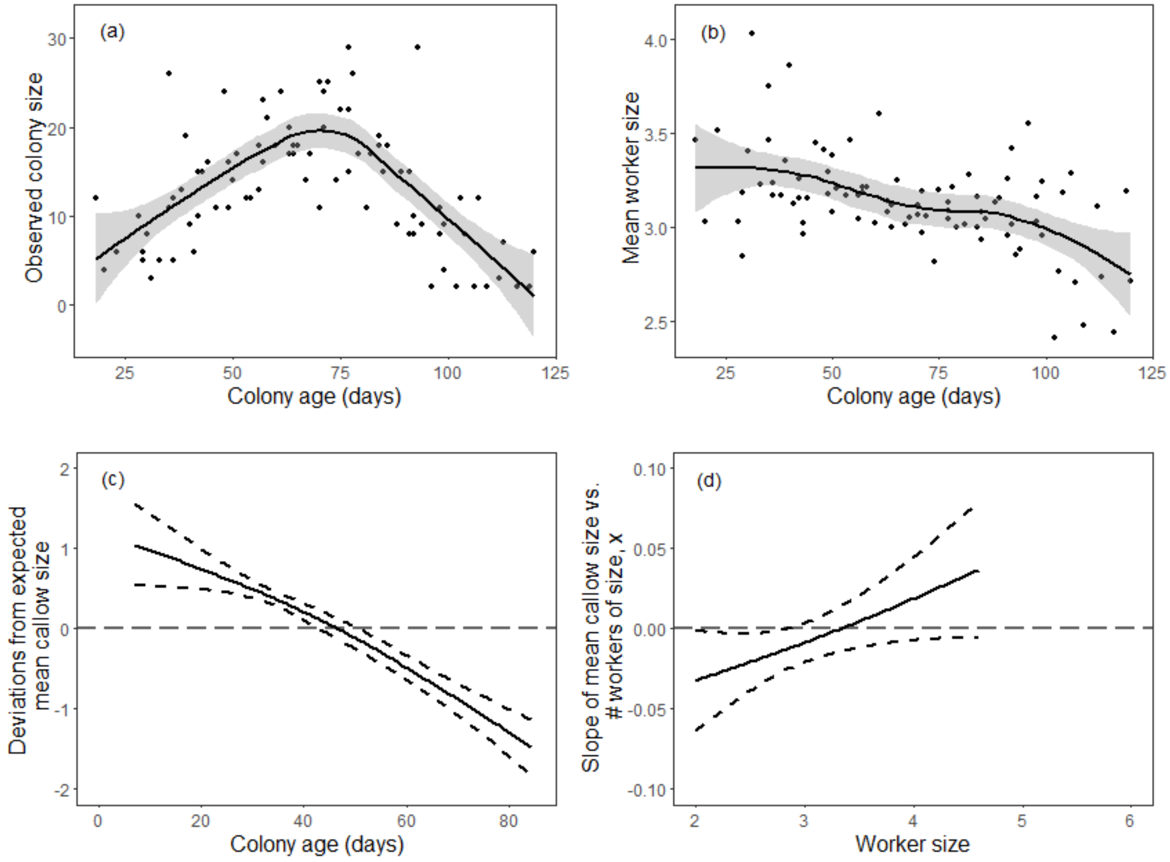


**Figure S4.7.** The (a) observed colony size (i.e. number of workers) and (b) mean worker size plotted against colony age in the low resource environment. Smooth components of generalized linear models evaluating the (c) deviations from expected mean callow size as a function of colony age and (d) the slope of mean callow size vs. # of workers as a function of worker size *x* for the low resource environment.

<u>High-low:</u> Neither colony age or worker size contribution had significant effects on mean callow size in the high-low resource environment. Therefore, we did not need to evaluate confounding effects for this vital rate.

<u>High:</u> Mean callow size decreased with increasing colony age (Fig S4.8c) and more larger workers (Fig S4.8d). Since worker size was the highest at younger colony ages (Fig. S4.8b), these smooth terms are not confounded.

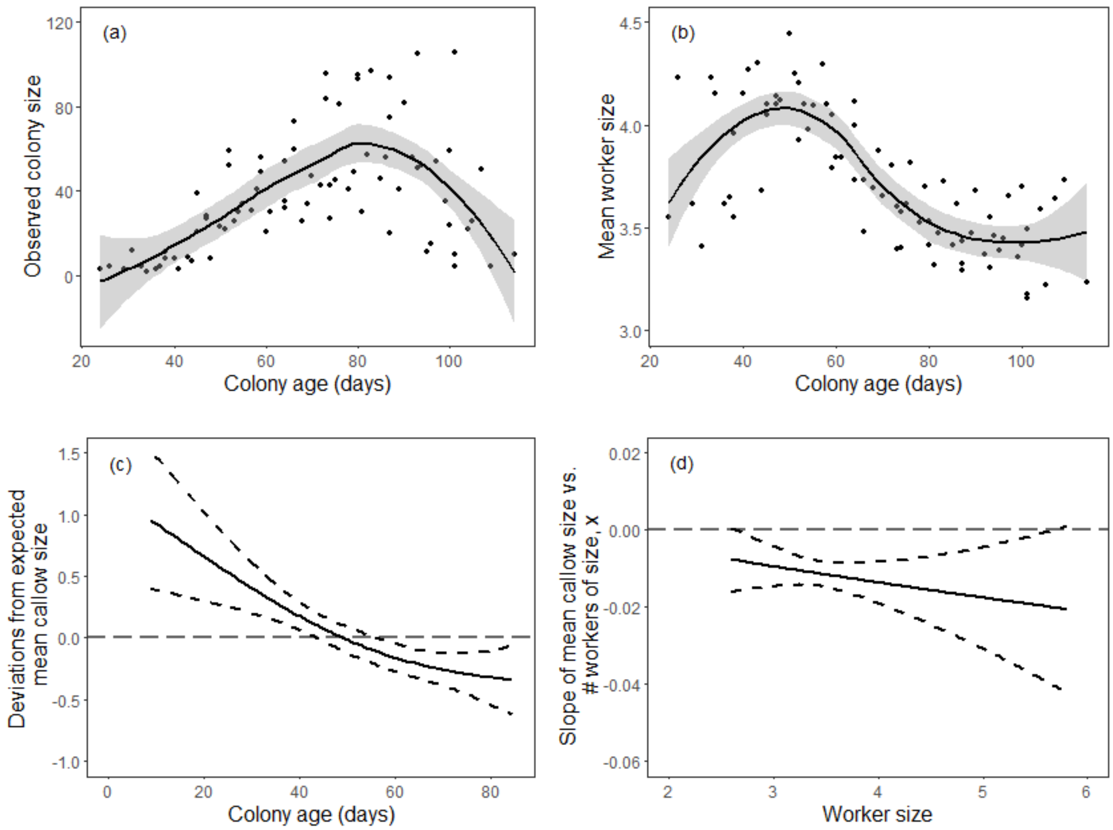


**Figure S4.8.** The (a) observed colony size (i.e. number of workers) and (b) mean worker size plotted against colony age in the high resource environment. Smooth components of generalized linear models evaluating the (c) deviations from expected mean callow size as a function of colony age and (d) the slope of mean callow size vs. # of workers as a function of worker size *x* for the high resource environment.

### 5. *Standard deviation in callow size*

Low: Neither colony age nor worker size contribution had significant effects on standard deviation in callow size in the low resource environment. Therefore, we did not need to evaluate confounding effects for this vital rate.

High-low: Colony age had no significant effects on standard deviation in callow size in the high-low resource environment, while worker size contribution had marginal effects. Therefore, we did not need to evaluate confounding effects for this vital rate.

High: Neither colony age nor worker size contribution had significant effects on standard deviation in callow size in the high resource environment. Therefore, we did not need to evaluate confounding effects for this vital rate.
